# ADOPT: intrinsic protein disorder prediction through deep bidirectional transformers

**DOI:** 10.1101/2022.05.25.493416

**Authors:** Istvan Redl, Carlo Fisicaro, Oliver Dutton, Falk Hoffmann, Louie Henderson, Benjamin M.J. Owens, Matthew Heberling, Emanuele Paci, Kamil Tamiola

## Abstract

Intrinsically disordered proteins (IDP) are important for a broad range of biological functions and are involved in many diseases. An understanding of intrinsic disorder is key to develop compounds that target IDPs. Experimental characterization of IDPs is hindered by the very fact that they are highly dynamic. Computational methods that predict disorder from the amino acid sequence have been proposed. Here, we present ADOPT, a new predictor of protein disorder. ADOPT is composed of a self-supervised encoder and a supervised disorder predictor. The former is based on a deep bidirectional transformer, which extracts dense residue level representations from Facebook’s Evolutionary Scale Modeling (ESM) library. The latter uses a database of NMR chemical shifts, constructed to ensure balanced amounts of disordered and ordered residues, as a training and test dataset for protein disorder. ADOPT predicts whether a protein or a specific region is disordered with better performance than the best existing predictors and faster than most other proposed methods (a few seconds per sequence). We identify the features which are relevant for the prediction performance and show that good performance can already gained with less than 100 features. ADOPT is available as a standalone package at https://github.com/PeptoneLtd/ADOPT.

## INTRODUCTION

Modern biochemistry is based on the assumption that proteins fold into a three dimensional structure, which defines their biological function.(1) However, in the past 20 years a novel class of biologically active polypeptides was identified that defies the structure-function paradigm. Intrinsically disordered proteins (IDPs) constitute a broad class that encompasses proteins which do not fold into a defined three dimensional structure, undergo transitions between multiple unstructured or partially structured conformations or fold upon binding. (2) IDPs play a fundamental role in biological processes such as cell signalling and regulation(3) and were implicated in numerous debilitating human disorders, including various cancers(4), diabetes(5), cardiovascular diseases(6) and neurodegenerative conditions like Alzheimer’s or Parkinson’s disease.(7) Consequently, IDPs are considered prime targets for drug development, yet very limited success of IDP-targeting compounds has been reported to date.(8) The development of therapeutic molecules that target full-length IDPs or unstructured regions of folded proteins (IDRs) is particularly challenging because unfolded regions are depleted in hydrophobic residues, which stabilize druggable protein regions. Instead, IDPs contain many charged and polar residues, whose repulsive interactions drive intrinsic protein disorder. Thus amino acid sequence composition of IDPs holds key to precise prediction and assessment of structural disorder in high-value and unmet medical targets. The rational design of IDP-targeting compounds is hindered by the lack of of a well-define structure and by the challenges posed by the experimental characterization of the broad conformational ensembles they populate. The big amount of potential IDP drug targets is presently not approachable with traditional structured based drug design tools and needs the development of new methods. Experimental techniques used to characterize intrinsic disorder range from X-ray crystallography, nuclear magnetic resonance spectroscopy (NMR), small-angle X-ray scattering, circular dichroism and Förster resonance energy transfer. The resolution of the information and nature itself of the disorder probed depends on the experimental technique used to characterize it. Nuclear Magnetic Resonance (NMR) chemical shifts depend uniquely on the local environment and provide a probe, at single amino acid level, of the structure and dynamics of IDPs.

Since the first disorder predictor was proposed in 1997(9), numerous protein disorder predictors have been developed. In general, they can be divided in four categories: physicochemical predictors, machine learning (ML) based predictors, template-based predictors and meta predictors.

Physicochemical property-based predictors(10, 11, 12, 13, 14, 15) utilise chemical features of amino acids, especially hydrophobicity and net charge, to predict if a residue belongs to an ordered or disordered region. Disorder predictions can be straightforwardly interpreted in terms of physical properties of underlying amino acid sequence. However, the accuracy and sensitivity of such predictors are limited to the features they use to differentiate between order and disorder. ML-based predictors(16, 17, 18, 19, 20, 21, 22, 23, 24, 25, 26, 27, 28, 29, 30) use positive and negative samples to distinguish between ordered and disordered regions. They can be used to search for the most important among many features. In comparison to physicochemical predictors, ML-based predictors are more flexible in their search of features, yet may suffer from poor ‘explainability’ of extracted features and their relevance for disorder. Template-based predictors(31, 32, 33) search for homologous structures (templates) and use these to distinguish between structured and unstructured regions in proteins. Their results can be easily interpreted, but homologous structures might not be found or their disorder prediction might not transferable to the query protein. Finally, meta predictors(33) aggregate the output of multiple other disorder prediction tools and report a composite disorder score. In general, this leads to a better prediction accuracy than individual predictors, but at the cost of an increased computational effort which in turn may hamper their effectiveness against proteome-size surveys. Despite the progress in disorder prediction in the last 25 years, a systematic assessment of the different predictors did not exist until a few years ago. Recently, two comparisons have been established. The Critical Assessment of Protein Intrinsic Disorder Prediction (CAID)(34) is a community-based blind test of state of the art prediction of intrinsically disordered regions based on 643 proteins from DisProt(35). DisProt, the most comprehensive database of disordered proteins provides manually curated annotations of currently about 2400 IDPs and IDRs of at least ten residues likely to be associated with a biological function. The data is based on multiple experimental measurements, but not all IDRs of a protein are contained in DisProt. Furthermore, the annotations in the DisProt database are binary, e.g., they do not report on the strength of disorder of a specific residue in the protein.

CheZoD is a small database(36) of 117 proteins known to contain disorder for which NMR chemical shifts are available from the BMRB database. (37) The CheZOD Z-score (referred to as Z-score below), based on secondary chemical shifts and defined in Ref (36), quantifies the degree of local disorder on a continuous scale. Secondary chemical shifts. e.g. differences in chemical shifts of nuclei between the actual structure and a random coil structure, are a precise indicator of local protein disorder.(38) It was demonstrated that the Z-score scale, besides being a reliable measure of disorder, also agrees well with other measures of disorder. The histogram of all Z-scores calculated for an expanded CheZOD database (39) containing 1325 proteins and constructed in a way to ensure balanced amounts of disordered and ordered residues fits a bimodal distribution; residues with Z-scores *<* 3.0 can be considered fully disordered, whereas 3.0 *<* Z *<* 8.0 corresponds to cases with fractional formation of local, ordered structure.

A systematic comparison of 43 predictors in CAID(34) and 27 predictors on Z-scores(39) showed that deep learning techniques clearly outperform physicochemical methods. For example, SPOT-disorder(40, 41) was the second best predictor in both competitions, slightly beaten by fIDPnn(42) in CAID and by ODiNPred(39) on Z-scores. All three predictors use deep neural networks to predict protein disorder.

The elucidation of information from protein sequences is one of the most formidable challenges in modern biology. A comparable task in Artificial Intelligence research is Natural Language Processing (NLP), in which ML and linguistic models are used to study how properties of language including semantics, phonetics, phonology, morphology, syntax, lexicon and pragmatics arise and interact. To accomplish this, NLP models must be able to find common patterns in variable length and non-linear correlations in language constituents as letters, words, and sentences.(43) The final goal of NLP is to find regularities of natural language and universal language creation rules.

Supervised learning, a rapidly emerging branch of artificial intelligence research, is of particularly high importance to computational biology where vast and well annotated data repositories are available. Its more sophisticated variant, self-supervised learning, enables Artificial Intelligence (AI) systems to learn from orders of magnitude more data than supervised ones, which is important for recognising and understanding patterns of more subtle and less common representations. Self-supervised neural models like the Transformer (44) have recently proven to be especially effective for these tasks, for example vastly improving language modeling and next sentence prediction in BERT (45) or generating high quality human-like text in GPT.(46)

Motivated by recent developments in self-supervised learning and its application to protein language models (47, 48), we developed a novel structural disorder predictor that benefits from the Transformer architecture. We demonstrate here, the relative benefit of ADOPT (Attention DisOrder PredicTor) software on the expanded version of the CheZOD database. ADOPT outperforms the state of the art tools, ODiNPred, SPOT-disorder, and MetaPredict v2, in a number of benchmarks. Finally, we provide an analysis of the residue level representations used in the development of ADOPT and discuss in detail the features that contribute to an accurate prediction of protein disorder from sequence alone.

## MATERIALS AND METHODS

ADOPT is composed of two blocks: a self-supervised encoder and a supervised disorder predictor. The encoder takes a protein input sequence and uses information from a large database of sequences to generate feature information for every residue in the sequence. The decoder uses this information and predicts a disorder score. We used Facebook’s Evolutionary Scale Modeling (ESM) library to extract dense amino acid residue level representations, which feed into the supervised machine learning based predictor. The ESM library exploits a set of deep Transformer encoder models (44, 45), which processes character sequences of amino acids as inputs. A high level representation of the ADOPT architecture is shown in Figure 1.

**Figure 1.**
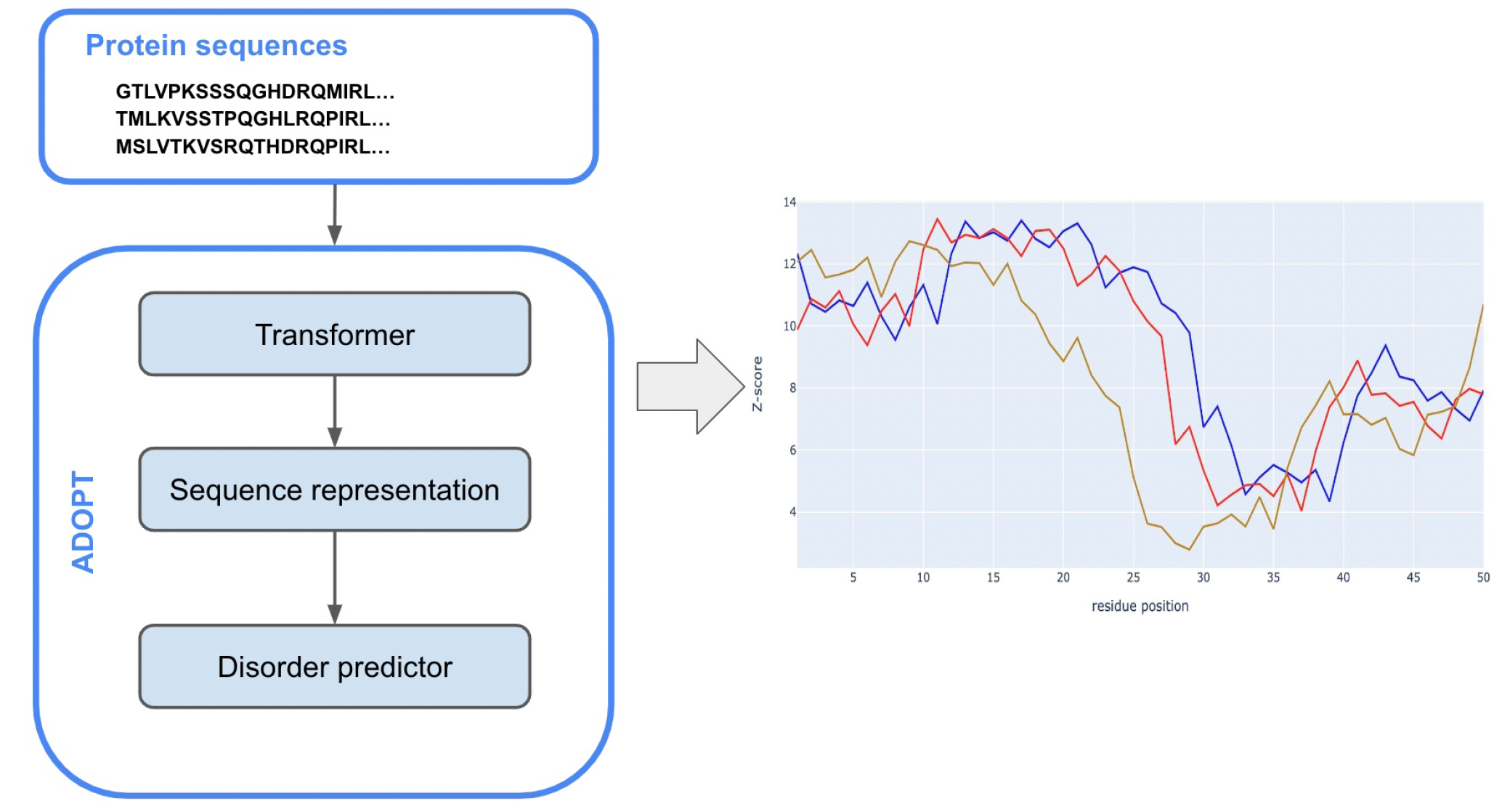
Schematic view of the ADOPT architecture. A protein sequence is fed into the Transformer encoder block which generates a residue level representation of the input. The embedding vector serves as an input to the disorder predictor block which predicts the level of disorder of each residue, given in terms of z-scores.

The encoder maps each element *x*_*i*_ ∈ *V* of an input sequence of symbol representations **x** =(*x*_1_,…,*x*_*n*_) to a dense vector **z**_*i*_ where 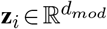. This means that the context of every residue within the input sequence is encoded in a vector with *d*_*mod*_ features. Here, *d*_*mod*_ ∈ ℕ^+^ is the *embedding dimension* which is set at training time, *n* ∈ ℕ^*^ is the length of the protein *p* represented by **x** and

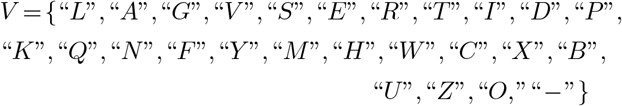

represents the vocabulary, where “*X*” stands for unknown amino acid, and “ − “ is the gap symbol in the event of a multi-sequence aligned input. Furthermore, “*B,U,Z,O*” are non-natural amino acids, and the remaining 20 are standard ones. Therefore a protein *p*, represented by the sequence vector **x** will be encoded on an embedding matrix 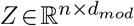 obtained stacking **z**_*i*_ for *i* = 1,2,…,*n*.

ADOPT computes residue level *Z-scores* (49), which quantify the degree of local disorder related to each residue, on a continuous scale, based on NMR secondary chemical shifts (50). The input of the disorder predictor is given by the dense representations (embeddings) *Z* of each protein sequence **x**, whereas the output is the predicted Z-score of each element *x*_*i*_ of **x**.

In order to strike a balance between readability and details, whilst keeping the paper self-contained, we provide more details on the transformer block, including the attention mechanism, see Supplementary Figure S1, in APPENDIX A in the Supplementary Material.

### Datasets

#### Pre-training datasets

Pre-training datasets were based on UniRef (51). The UniRef50 and the UniRef90 were extracted from UniParc (52) by clustering at 50% and 90% sequence identity, respectively. The UR90/S dataset represents a high-diversity and sparse subset of UniRef90 dated March 2020 and contains 98 million proteins, while the UR50/S dataset represents a high-diversity and sparse subset of UniRef50 representative sequences from March 2018 with 27.1 million proteins. The MSA-UR50 dataset was generated with Multiple Sequence Alignment (MSA) to each UniRef50 sequence by searching the UniClust30 (53) database in October 2017, with HHblits 3.1.0 (54) and contained 26 million MSAs. Default settings were used for HHblits except for the the number of search iterations (`-n`), which was set to 3. Sequences longer than 1024 residues were removed and the average depth of the MSAs was 1192.

#### Disorder prediction datasets

The disorder prediction datasets were the CheZoD “1325” and the CheZoD “117” databases (39) containing 1325 and 117 sequences, respectively, together with their residue level Z-scores. Note that, throughout this paper we refer to the “1325” and “117” sets as *base* and *validation* sets, respectively.

Although, Z-scores can be transformed into probabilities of disorder that report on the likelihood of a residue being disordered, we decided to adhere to Z-scores primarily reported in ODiNPred (39).

### Pre-trained Transformer Models

ADOPT utilizes three different pre-trained Transformers: *ESM-1b, ESM-1v* and *ESM-MSA*. The architectures of *ESM-1b* and *ESM-1v* were described above and pre-trained on UR50/S and UR90/S, respectively. A complete review of *ESM-MSA*, architecture is given in (55). The aforementioned Transformer was pre-trained on MSA-UR50; see Table 1.

**Table 1.** Hyperparameters, dataset and number of parameters related to each ESM model employed.

| Transformer | Dataset | $l$ | $d_{mod}$ | $h$ | Params |
| --- | --- | --- | --- | --- | --- |
| ESM-1v | UR90/S | 33 | 1280 | 20 | 650 M |
| ESM-1b | UR50/S | 33 | 1280 | 20 | 650 M |
| ESM-MSA | MSA-UR50 | 12 | 768 | 12 | 100 M |

### Disorder predictors

We built four different variants of ADOPT, which differed in the underlying ESM model. We used *ESM-1v, ESM-1b, ESM-MSA* and *ESM-combined*, where the latter one referred to the concatenation of the representations given by *ESM-1v* and *ESM-1b*. The input dimension *d*_*mod*_, which was equivalent to the dimension of the residue level representation, was dependent on the ESM transformer used (Table 1), for example *d*_*mod*_ = 2560 for *ESM-combined* and *d*_*mod*_ = 1280 for *ESM-1b*. Note that *ESM-combined* is an artificial concept and just a label that not represent a standalone transformer model. In fact, it is only a reference to the concatenated outputs extracted from the two pre-trained transformers *ESM-1v* and *ESM-1b*.

Not all four predictors were tested in every benchmark exercise due to the fact that some of them, e.g. *ESM-1v* and *ESM-1b*, have similar performances. We indicate in each result discussed which predictor is used.

To predict Z-scores using the residue level representations, we used a simple Lasso regression model with shrinkage parameter *λ* = 0.0001, which is considered a well known, standard technique, see, e.g. chapter 3 in (56). The optimal value for *λ* was found experimentally by a search across of the interval [0.0001,0.5].

In the binary case, i.e. predicting *order/disorder* only, we used a simple logistic regression with a *L*_2_ penalty term, for details, see chapter 4 of the same reference (56).

As a benchmark with AlphaFold2, we correlated rolling averages of pLDDT and SASA, defined in Benchmark on order/disorder classification subsection, with Z-scores. In both metrics the window lengths were ranging from 5 to 30 in steps of 5. In the three cases of sequences containing unknown amino acids the unknown amino acid was replaced with glycine. SASA was calculated using FreeSASA ^1^ 2.0.3 with default parameters, with the total relative per residue used for correlations.

### Supervised training and evaluation

Our predictors were trained on the *base* data set reduced by 89 sequences, that were found to display an overlap between the base and the validation set, where overlap was defined as minimum 50% sequence identity, i.e. sequences that shared at least 50% sequence similarities. These were identified with the tool `MMSeqs2`; see (57) and https://github.com/soedinglab/MMseqs2. Hence the final training set consisted of 1236 sequences. We used the validation set containing 117 sequences as the test set. The performance of our regression models was measured by Spearman correlation *ρ*_Spearman_ between predicted and experimental Z-scores. To further assess model performance, we also provided mean absolute errors (MAE), Average prediction errors. Where relevant, we also added *p*-values. However, given that these were consistently 0.0, i.e. detecting e.g. correlation falsely had zero probability, we omitted from most Tables.

Following closely the cross validation settings used in (39), we performed 10-fold cross-validations (CV), separately, only using the base data set, i.e. 1325 sequences. We did not remove any sequences for the (CV) exercises, i.e. the total number of sequences was 1325. As per standard practice, in a fold 9*/*10-th of the base data set was used as the training set, where model was fitted, and the remaining 1*/*10-th as test set. We used randomised 10-fold (CV), that is, the folds were selected randomly. A new regression was trained for each fold and we report the average correlation *ρ*_Spearman_, average MAE, and average prediction errors, together with standard deviations, taken across the folds. For the regression task, we performed two cross-validation exercises; one based on *residue level* and another on *sequence level* fold selection. In the former case, whether a residue was in the training or test set was decided on amino acid level. The latter meant that training and test sets were assembled by sequences, i.e. all residues in a sequence were either in the training or in the test set.

In the order/disorder classification tasks, we used Receiver Operating Characteristic Curve (ROC AUC) and Matthews Correlation Coefficients (MCC) as standard evaluation metrics. Furthermore, we provided precision scores, whereby ordered residues were treated as ‘positives’ (Z-scores higher than 8.0). Our classifiers were evaluated in two settings, a residue level 10-fold CV on the base set, and a training using the reduced base set and testing on the validation set.

### Feature selection

Two methods were used to identify a subset of relevant coordinates or features of the residue level representation vectors, as shown in Figure 5. Recall that we used Lasso regression as our disorder predictor.

In *naive selection*, we selected only those coordinates, whose coefficients were above a certain threshold in terms of their absolute value, that is, the subset *S* ⊂ {1,2,…,1280} was given by

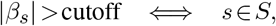

where *β*_*i*_ for *i* ∈ {1,2,…,1280} are the Lasso coefficients for the fixed shrinkage parameter *λ* = 0.0001. We fitted a new linear regression (Lasso with the same parameter *λ* = 0.0001) on the reduced set of features and reported the correlation *ρ*_*Spearman*_ between predicted and actual Z-scores on the test set. We repeated this procedure for 10 different *cutoff* values, equally spaced ranging from 0.5 to 0.2. This selection method is called naive, as it throws away coordinates whose coefficients are below the cutoff in absolute terms, which may still be relevant, and the set of selected features may contain a fair amount of noise. ‘ As an alternative method we used *stability selection* to identify relevant subsets of the representation vector coordinates. We defined 30 different shrinkage parameters *λ*’s. These were equally spaced points of the interval [5 × 10^−5^,0.1], endpoints included. For each *λ* we carried out the following procedure. From the initial test set, we chose a random subset, whose size was half of the original one. We fitted a Lasso regression and recorded the features, whose coefficients were non-zero. We repeated this 500 times and for each vector coordinate *i* ∈ {1,2,…,1280} we estimated the selection probability Π_*i*_(*λ*) by the number of times the coordinate got selected divided by 500.

In order to arrive to the relevant subsets, we applied cutoff values at two different stages. First, we put a threshold *c*_*p*_ on the selection probabilities. Second, we counted for how many *λ* values this particular threshold *c*_*p*_ has been exceeded. We refer to the latter quantity as frequency cutoff and denote it by *c*_*f*_. We used the following values

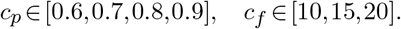

As an example, for *c*_*p*_ = 0.6 and *c*_*f*_ = 15, a representation vector coordinate *i* was deemed relevant if the selection probability Π_*i*_(*λ*) *>* 0.6 for at least 15 different *λ* values. The estimated probabilities Π_*i*_(*λ*) as a function of *λ* ∈ Λ can be depicted as paths. We use the term *stability paths* to describe those that pass the aforementioned two thresholds *c*_*p*_ and *c*_*f*_, as shown in Figure 6. For more details and discussions on stability selection in general we refer to (58).

### Implementation

ADOPT is available as a command-line utility at https://github.com/PeptoneLtd/ADOPT. The software uses Facebook’s ESM^2^ 0.4 for the pre-training phase, Sklearn ^3^ 1.0 for the disorder predictor, BertViz^4^ 1.2 for the multi-head attention visualisation, HHblits^5^ 3.1.0 for extracting the MSAs and ONNX^6^ 1.10.2 for the inference phase. The embedding representations were extracted in bulk with the ESM command-line utility specifying the `--include per tok` option from both CheZod “1325” and CheZod “117” FASTA files. The embedding vectors extraction and tool development (training and inference) were performed on a single Nvidia DGX A100.

## RESULTS

### Disorder predictor developed using the CheZoD data sets

In order to compare ADOPT to the current state-of-the-art disorder predictor ODiNPred, we adopted the benchmarking protocol found in (39). The CheZoD database was split into two parts, a *base* and a *validation* set, containing 1325 and 117 sequences, respectively, with their residue-level Z-scores, which are NMR-based measures of disorder. We used Z-scores to compute the probability of disorder ℙ_disorder_ at residual level. A probability close to 1 indicates that the residue is likely disordered. Throughout this paper we are only considering Z-scores as a measure of disorder and omit any further quantification of probabilities of disorder. Furthermore, for classification tasks residues with Z-scores below/above 8 were assumed to be disordered/ordered, respectively.

### Training on the base set and performance evaluation on the validation set

We evaluated the variants of ADOPT on four different representations obtained from ESM-transformers: *ESM-1b, ESM-1v, ESM-MSA* and also an *ESM-combined* that used the concatenated representations from *ESM-1b* and *ESM-1v*. Note, that the choice of the transformer model changes the input of our disorder predictor and therefore influences its performance. The reduced base and validation sets consisted of 1236 and 117 sequences, respectively. Here, sequences which are part of both sets of the CheZoD database were removed from the *base* set. We found that disorder predictors had a tendency to overestimate the experimental Z-scores, which was a phenomenon universally observed across the whole spectrum of predictors (39). On the validation set ODiNPred e.g. displayed an average prediction error of − 2.7226, meaning that it typically predicted higher Z-scores, and a mean absolute error (MAE) of 3.7348. Although, our ESM-based predictors also overestimated Z-scores, their prediction errors were considerably lower, e.g. for *ESM-1b* the average prediction error and MAE were − 2.1124 and 3.3791, respectively. Spearman correlation coefficients *ρ*_Spearman_ between the predicted and actual Z-scores were used to benchmark performance against ODiNPred. Our ESM-transformer based predictors showed the following Spearman correlation coefficients between predicted Z-scores and experimental Z-scores: 0.689 (*ESM-1b*), 0.681 (*ESM-1v*), 0.615 (*ESM-MSA*) and 0.694 (*ESM-combined*); compared to the reported Spearman correlation coefficient of 0.649 for ODinPred and 0.64 for SPOT-disorder given the same prediction task (39); see Table 2.^7^ We also performed the same benchmark test on a recent disorder predictor, Metapredict v2(59). Metapredict V2 was generated by training a neural network on the disorder scores of a hybrid version of the original Metapredict disorder predictor(60) and predicted pLDDT scores. While the original predictor was highly competitive in the last CAID competition, the successor clearly outperforms the performance ot the original predictor with faster execution time. The Spearman correlation coefficient between predicted disorder scores from Metapredict V2 and experimentally derived scores was 0.6581, slightly better than ODinPred and SPOT-disorder, but worse than ADOPT. In summary, ADOPT predicted protein disorder more accurately than current state-of-the-art predictors and the overestimation of experimental Z scores observed across all the tested disordered predictors indicates that these ML-based algorithms display high sensitivity to sequence patterns present in ordered regions.

**Table 2.** Spearman correlations between actual Z-scores and predicted pLDDT_5_ scores along with actual Z-scores and predicted SASA_15_ scores, obtained by AlphaFold2. Spearman correlations between actual and predicted Z-scores, average prediction error (actual - predicted) and mean absolute error (MAE) obtained by ODiNPred and our ESM-transformer based predictors. These correlations have been collected for the task linked to the model evaluated on the *validation set* consisting of 117 sequences. Given that the *p*-values, in all cases were 0.0, we omitted them below.

| Disorder predictor | Spearman<br>corr. coeff.<br>($\rho_{\text{Spearman}}$) | Avg.<br>prediction<br>error | MAE |
| --- | --- | --- | --- |
| AlphaFold2, pLDDT<sub>5</sub> | 0.5548 |  |  |
| AlphaFold2, SASA<sub>15</sub> | 0.6299 |  |  |
| MetaPredict v2 | 0.6581 |  |  |
| ODINPred | 0.649 | $-2.7226 \pm 3.95$ | $3.73 \pm 3.01$ |
| ESM-1v | 0.6808 | $-2.2747 \pm 3.94$ | $3.5057 \pm 2.89$ |
| ESM-1b | 0.6886 | $-2.1124 \pm 3.84$ | $3.3791 \pm 2.79$ |
| ESM-MSA | 0.6147 | $-2.8588 \pm 4.11$ | $3.9563 \pm 3.07$ |
| ESM-combined | **0.6938** | **$-2.0917 \pm 3.89$** | **$3.3894 \pm 2.83$** |

The prediction of local disorder in a residue is dependent on the neighborhood of this residue. In order to estimate if our predictor is able to correctly identify disordered regions, we computed the disordered regions prediction recall score i.e. the fraction of correctly predicted disordered regions from all disordered regions in the dataset. First, the Z-score related to each residue in all proteins of the *validation* set has been predicted with the ESM-1b model as upstream task. After converting the Z-scores of both, the ground truth and the predicted ones, into a binary class, i.e., *Z <* 8 for a residue belonging to a disordered region and *Z* ≥ 8 for a residue in a ordered region(39), we identify for each sequence the related disordered regions defined as those areas where the number of consecutive disordered residues is equal or greater than 3 in the ground truth and 70% of the residues in those areas are disordered in the prediction. The recall computed over the binary class of disordered regions is 66%. This means that the ADOPT disorder predictor trained on the ESM-1b transformer is able to correctly find most disordered regions.

Furthermore, we test the robustness of our methods. Therefore we introduce two small changes in the sequence and measure the change in disorder prediction. First, we predict how the termini of a protein sequence which are less structured and more flexible influence our predictions. We trim between one and ten residues at both termini of each sequence and measure the difference between predicted and actual Z scores for the rest of the protein. Second, we randomly mutated up to 10 residues in the *validation* dataset. In both cases, we report the differences in terms of mean absolute error and Spearman correlation computed on the related ground truth and predicted Z-scores as well as recall score computed over disordered regions binary class. The results in Figure 2 show that the accuracy of ADOPT is not influenced by the inclusion of random residues and changes in the number of termini residues showing that ADOPT is robust upon small changes in the given input protein sequence.

**Figure 2.**
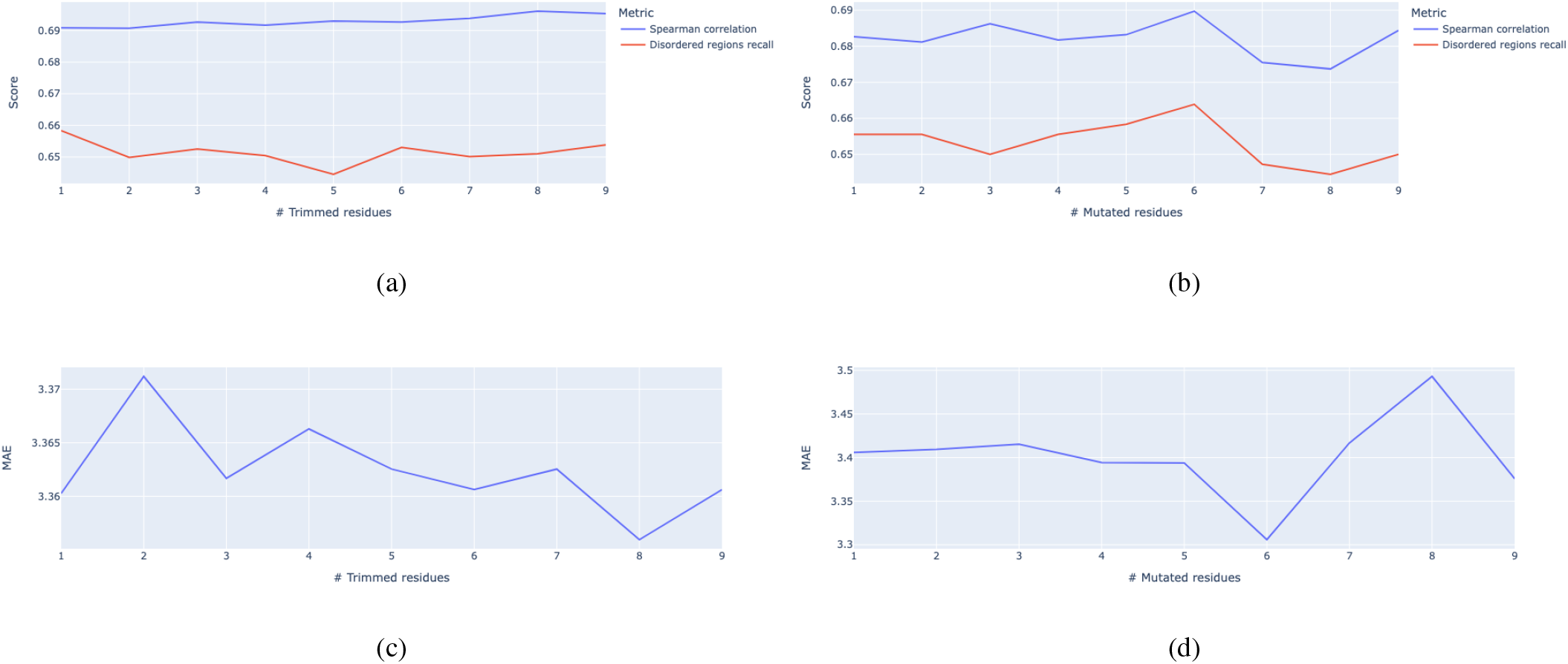
Reliability study on ADOPT predictions. Spearman correlations and disordered regions prediction recall computed on trimmed and randomly mutated sequences are shown in Figure 2a and Figure 2b, respectively. Mean Absolute Error (MAE) computed on trimmed and randomly mutated sequences are shown in Figure 2c and Figure 2d, respectively

In order to understand the origin of improvement observed in ADOPT, we compared the performance of one of our ESM-transformer based predictors *ESM-1b* against ODiNPred by visualising the contour plots of the 2D density of actual vs. predicted Z-scores, see Figure 3. Here, the reference lines indicate the idealized case, where predicted values are equal to the actual values. The more probability mass lies closer to these reference lines, the better the accuracy of the predictor is. Figure 3 demonstrates that both predictors identify correctly the bimodal Z score distribution of ordered and disordered regions. This distribution has been chosen in the development of the CheZoD database in order to have a good mixture of both protein classes. The average mass is above the reference line in both regions and for both predictors, which shows that the bias to ordered regions is not limited to a specific strength of protein disorder. The contour lines are tighter around the reference line for the ESM-transformer based predictor (on the left), than for ODiNPred (on the right). The data for ODiNPred was obtained via the publicly available web service https://st-protein.chem.au.dk/ odinpred. Note that using these predicted Z-scores a slightly higher Spearman correlation coefficient of 0.671 was found for ODiNPred, than 0.649 as reported in the paper (39). The result shows that the ESM-transformer produces less outliers in comparison to ODiNPred. While the prediction of the ESM-transformer is far from perfect, the probability of returning a bad Z score prediction, defined by a difference of Δ*Z*=5 from the experimental Z score, is significantly lower than for ODiNPred, see Figure 3.

**Figure 3.**
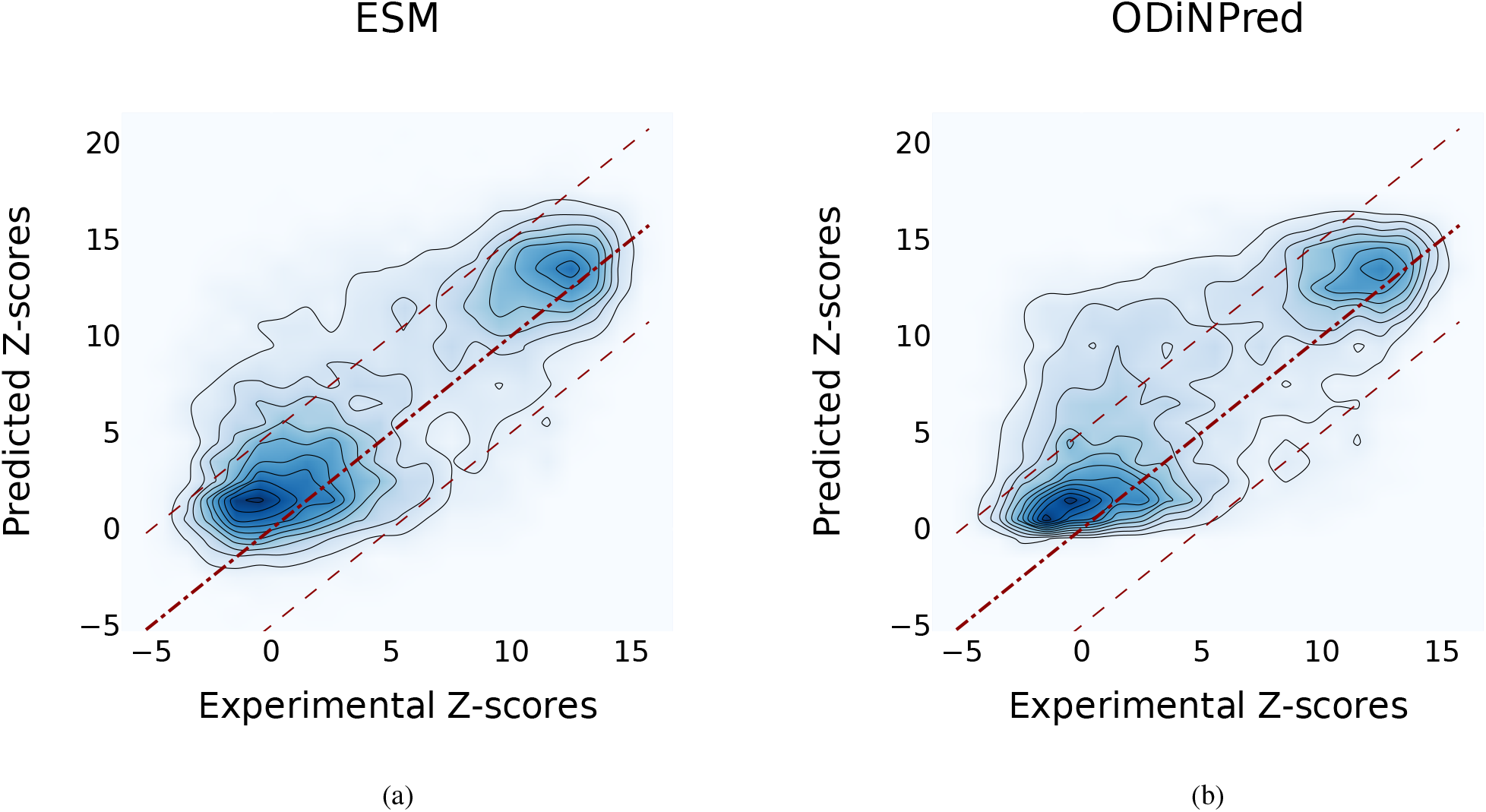
Contour plots showing the density levels of experimental vs. predicted Z-scores for the validation set containing 117 sequences; Figure 3a shows results ESM-transformer based predictor (here *ESM-1b* was used); Figure 3b shows the results obtained with ODiNPred. Diagonal reference lines indicate the idealized “perfect” prediction, i.e. predicted values are equal to the experimental values. As the probability mass around the reference line is tighter for the ESM based predictor on the left, its accuracy is better than that of ODiNPred. This is further supported by the average prediction error and MAE for the two predictors, see Table 2. Z-scores predicted by ODiNPred were retrieved via the publicly available web service referenced in (39). Note that using these predicted Z-scores a slightly higher Spearman correlation of 0.671 was found, rather than 0.649 as reported in (39). Recall that the Spearman correlation for *ESM-1b* was 0.6886.

Our amino acid level analysis reveals further details about the biophysical origin of the prediction accuracy. We compared two of our proposed disorder predictors, *ESM-1b* and *ESM-combined*, to ODiNPred (Figure 4). Keeping the same setting as already described above, i.e. *ρ*_Spearman_ between the predicted and experimental Z-scores for the validation set, we observed that the ESM-transformer based predictors displayed better than or similar correlation levels to that of ODiNPred for most amino acids. Some notable exceptions were Histidine (H) and Methionine (M). In these cases, ODiNPred was significantly weaker with correlation coefficients *ρ*_Spearman_ = 0.5843 and *ρ*_Spearman_ = 0.5607 compared to e.g. *ESM-1b* with *ρ*_Spearman_ = 0.6525 and *ρ*_Spearman_ = 0.6685 for (H) and (M), respectively. Curiously, all three predictors had considerable difficulties with glycine (G), cysteine (C) and proline (P). In fact, for Glycine, ODiNPred’s *ρ*_Spearman_ = 0.6102 outperformed both ESM-transformer based predictors by about 0.03; see Figure 4. The distribution of Z scores for these amino acids (Supplementary Figures S2 and S3) demonstrates that especially the Z scores of ordered glycine and proline residues are shifted to higher Z scores in the predictions of *ESM-1b* and ODinPred in comparison to the actual values. While this trend is observed for all residues, it has a more pronounced effect on the correlation coefficient of glycine and proline because these amino acids are underrepresented in ordered regions. The distribution of Z scores from cysteine (see Supplementary Figure S3) is clearly different from the distribution of the other amino acids with a higher relative amount of cysteines with intermediate Z scores around 8. ODinPred and *ESM-1b* however predict a much higher amount of ordered cysteine residues. Glycine, cysteine and proline are structurally different from the other 17 naturally occurring amino acids. Proline is the only amino acid with a secondary *α* amino group. Glycine is the smallest and most flexible amino acid, i.e., the only residue that has no chiral C_*α*_ atom. Both amino acids are commonly referred to as *α* helix breakers and thus are crucial for a correct prediction of disordered protein regions.(61) Cysteine has a thiol group and is therefore the only residue that is able to form disulfide bonds within the protein over long distances in sequence. Disulfide bonds are particularly important for the stability of proteins.(62) The forming or breaking of disulfide bonds depends on the oxidation state of the protein. This renders a correct prediction of its disorder state particularly challenging because the data we use to learn from and train on does not explicitly include the effect of oxidation. Moreover, the prediction of protein disorder remains especially challenging for amino acids which are important for the breaking of ordered regions. The newly developed ESM-transformer improves the prediction of disordered segments for 19 out of 20 amino acids. This shows that the improvement of the ESM-transformer is not limited to amino-acid specific biophysical properties.

**Figure 4.**
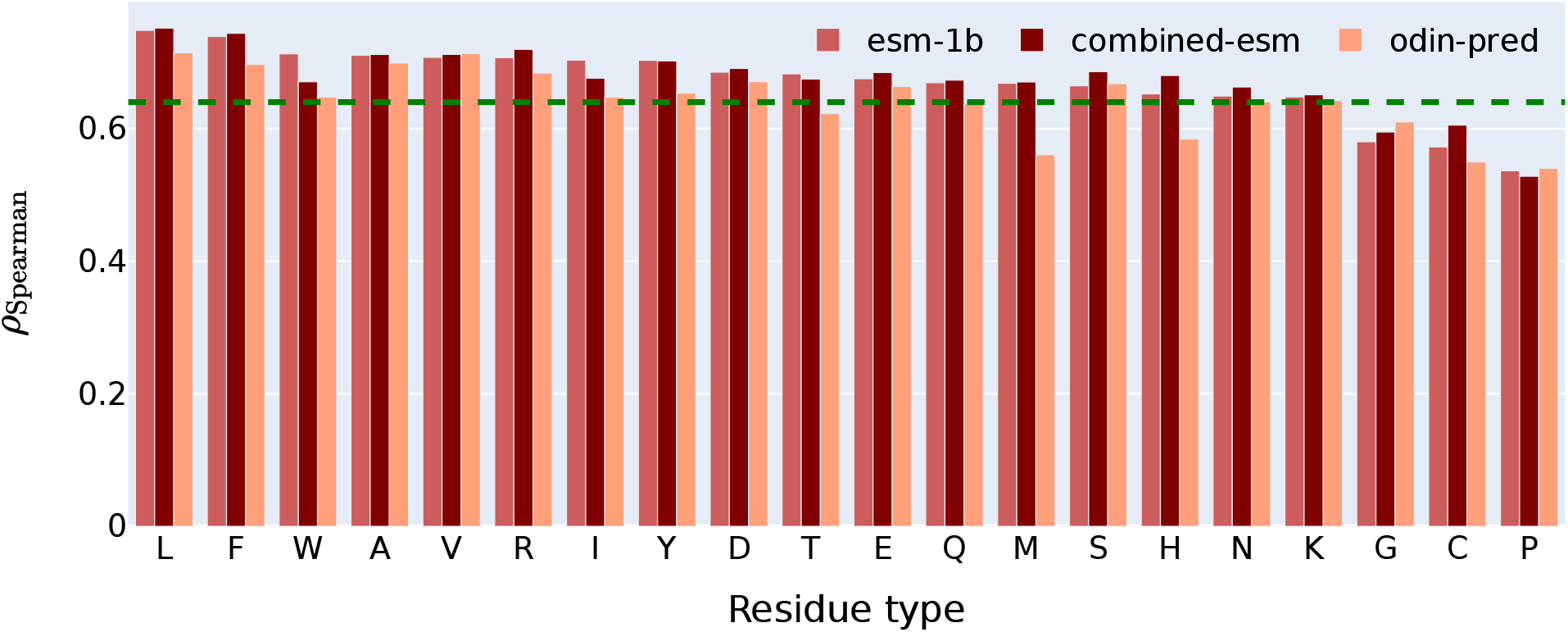
Residue level comparison of Spearman correlations between actual vs. predicted Z-scores for the validation set containing 117 sequences. Two of the ESM-transformer based predictors were used, *ESM-1b* and *ESM-combined*, which is based on the concatenated representations of *ESM-1v* and *ESM-1b*. The green reference line indicates the overall Spearman correlation of 0.649 obtained by ODiNPred as reported in (39).

### Performance of the ESM-transformer based disorder predictors assessed by 10-fold cross validations

To gain further insights into the performance of our predictors, we carried out two different 10-fold cross validations (CV) following the protocol proposed in (39); a detailed description of these tests is provided in the MATERIALS AND METHODS section. Cross validation gives information how the predictor generalizes to an independent dataset by resampling the validation set in different portions to avoid overfitting or selection bias. In the *residue level* 10*-fold CV* the Spearman correlations averaged across the 10 folds were as follows: 0.7576 (*ESM-1b*), 0.7577 (*ESM-1v*), 0.7322 (*ESM-MSA*). The same correlations for the *sequence level* 10*-fold CV* showed slightly lower values: 0.7342 (*ESM-1b*), 0.7324 (*ESM-1v*), 0.7160 (*ESM-MSA*). Under the conservative assumption that ODiNPred used *sequence level* fold selection, our ESM-transformer based predictors outperform ODiNPred by a considerable margin, which reported a Spearman correlation of 0.6904 for a similar 10-fold CV carried out on the base data set, see (39). These results are summarized in Table 3 (residue level) and Table 4 (sequence level) and show that our predictor does not suffer from selection bias, e.g., from the specific choice of the *validation set*, but rather gives better performance than ODinPred independent of the constitution of the *validation set*.

**Table 3.** Spearman correlations, Mean Absolute Error (MAE) and Average prediction error between predicted and actual Z-scores obtained by ODiNPred and our ESM-transformer based predictors for the *residue* level 10-fold cross validations on the *base set* consisted of 1325 sequences only.

| Disorder predictor | *residue level 10-fold CV* |  | Avg. prediction error |
| --- | --- | --- | --- |
| | $\rho_{\text{Spearman}}$ | MAE | |
| ODinPred | — | — | — |
| ESM-1v | **0.7577** | **2.3057** ± 0.0162 | **0.0002** ± 0.032 |
| ESM-1b | 0.7576 | 2.3079 ± 0.0147 | −0.0003 ± 0.030 |
| 5ESM-MSA | 0.7322 | 2.6457 ± 0.0188 | 0.0003 ± 0.042 |

**Table 4.** Spearman correlations, Mean Absolute Error (MAE) and Average prediction error between predicted and actual Z-scores obtained by ODiNPred and our ESM-transformer based predictors for the *sequence* level 10-fold cross validations on the *base set* consisted of 1325 sequences only.

| Disorder predictor | *sequence* level 10-fold CV |  | Avg. prediction error |
| --- | --- | --- | --- |
| | $\rho_{\text{Spearman}}$ | MAE | |
| ODINPred | — | — | — |
| ESM-1v | 0.7329 | $2.4477 \pm 0.0885$ | $0.004 \pm 0.239$ |
| ESM-1b | **0.7336** | **$2.4294 \pm 0.063$** | **$-0.007 \pm 0.187$** |
| ESM-MSA | 0.7153 | $2.7485 \pm 0.186$ | $-0.021 \pm 0.274$ |

### Benchmark on order/disorder classification

Experimental Z-scores from the sequences in the CheZoD database exhibit traits of a bimodal distribution, as shown in Figure 3. In (39), a mixture of two skew-normal distributions was fitted to the distribution of observed Z-scores with a reasonably high accuracy quantified by a Hellinger distance of 0.00367. This means that the distribution of Z scores of all sequences in the CheZoD database can be approximated by two distributions which are separated in Z score. This motivates a binary classification task, where a Z-score below/above 8 is interpreted as disorder/order, respectively. To remain aligned with (39), a 10-fold CV was performed on the base set. Accuracy was measured using standard metrics: the Area Under the Receiver Operating Characteristic Curve (ROC AUC) and the Matthews Correlation Coefficients (MCC) that measure dependency between binary outcomes. Here, a simple Logistic Regression was used on the representations extracted from the ESM-transformers. The ROC AUC / MCC averaged across the 10-folds were as follows: 0.9644*/*0.8044 (*ESM-1b*), 0.9643*/*0.8047 (*ESM-1v*), 0.9488*/*0.7729 (*ESM-MSA*). These results clearly exceed the values of *ROCAUC* = 0.914 and *MCC* = 0.690 ODiNPred reported for the same task in (39), see Table 5. In the aforementioned Table we also present precision scores for ESM-based predictors, for ODiNPred this metric was not reported in (39). Given that we labeled ordered residues with 1 (i.e. treated as ‘positives’), precision in this case means the percentage of the correctly predicted ordered residues. Our predictors are able to predict about 90% of ordered residues.

**Table 5.** ROC AUC, MCC and Precision by ODiNPred and our ESM-transformer based predictors for order/disorder classification given by the *residue* level 10-fold cross validations on the *base set* consisted of 1325 sequences.

| Disorder predictor | residue level 10-fold CV |  |  |
| --- | --- | --- | --- |
|  | ROC AUC | MCC | Precision |
| ODINPred | 0.914 | 0.690 | — |
| ESM-1v | 0.9643 | **0.8047** | 0.9078±0.002 |
| ESM-1b | **0.9644** | 0.8044 | **0.9081**±0.002 |
| ESM-MSA | 0.9488 | 0.7729 | 0.8937±0.002 |

In order to support these results further, a similar Logistic Regression was fitted to the base data set reduced by the overlap with the validation set. As described above, this data set contains 1240 sequences. The performance of the ESM-transformer based (here *ESM-1b*) order/disorder classifier was evaluated on the validation set containing 117 sequences. The results of *ROCAUC* = 0.8969 and *MCC* = 0.6185 obtained on this task can still be considered strong. Our analysis demonstrates that the ESM-transformer does not only predict better Z scores than ODiNPred, but also performs better on a binary classification of ordered and disordered regions, a topic which is very relevant in biology. This result is further reinforced by the lower mass of the ESM transformer in comparison to ODiNPred in the intermediate region with Z scores around 8 in Figure 3.

Additionally, it is worth mentioning that there are well known frameworks, e.g. CASP10, for disorder prediction assessment. As described in (39), the datasets in CASP10 were heavily imbalanced and less than 10% of the residues were disordered. To put this in context, in the overall CheZOD dataset, this percentage was 36.3%. The authors in (39) make two further observations. ODiNPred was not the best performing predictor in the CASP10 dataset, which may be due to the over-representation of ordered residues. Second, other disorder predictors overall performed much better on the CheZOD dataset.

The scientific community has shown great interest in AlphaFold2. Recent work has demonstrated that the predicted local distance difference test (pLDDT) (63), a per-residue metric of local confidence of the prediction, is correlated with disorder, outperforming IUPred2 (64), SPOT-disorder2(65) and other disorder predictors(66, 67). Further work has proved that the solvent accessible surface area (SASA) per-residue of models produced by AlphaFold2 shows better correlation to protein disorder than the pLDDT and that using a smoothing window improves accuracy further (64). To benchmark performance, AlphaFold2 structural models were predicted for the validation set of 117 proteins using the full database. For the rolling average of pLDDT and of SASA we observed that the largest correlations (0.5546 and 0.6299) were given by window sizes 5 and 15, respectively, as reported in Table 2. We note that predicting the structure from pre-calculated features derived from the MSAs took approximately 40 hours on one NVIDIA A100 GPU in total.

### A fast inference tool

From an end user perspective obtaining accurate disorder predictions quickly and easily is important. In this aspect ADOPT offers unmatched performance. We demonstrate this through comparisons with two other tools. ODiNPred currently is available through the web service https://st-protein.chem.au.dk/odinpred. Querying 117 proteins took overnight. We also made an attempt to use SPOT-Disorder2 https://sparks-lab.org/server/spot-disorder2/, which is considered at least as accurate as ODiNPred. However, a maximum of 10 sequences can be submitted at a time and the output is returned in 12 hours in the best case. In contrast, ADOPT, with a single command line, takes a few seconds to produce disorder predictions for 117 sequences and there is no cap on the number of sequences used in inference.

To prove that the ADOPT performance are the best in class not only in terms of disorder prediction but even in terms of inference time, we used the ADOPT to predict the Z-scores of the 20,600 unique protein coding genes from the reference *human proteome*^8^ which encode for 79,038 protein transcript variants. The protein sequences longer than 1024 units were split in two sub-sequences, due to the ESM context window length constraint mentioned above. The whole procedure takes roughly *15 minutes* to be completed on a single NVIDIA DGX A100, which demonstrates that the ADOPT inference phase is three orders of magnitude faster than that of OdinPred and AlphaFold2. However, this comparison is limited taking advantage of the fact that ADOPT runs as a fast command line tool while ODinPred and SPOT-Disorder2 have been used as a web service. To give a fairer comparison, we repeated the speed benchmark with Metapredict v2 on 1 CPU core, which reflects the standard usage of an end user of this predictor. Running the *human proteome* took 15 and 46 minutes on an Apple M1 chip and on a Colab instance, respectively, and is therefore of comparable speed to ADOPT using a GPU. Note, that Metapredict v2 has been generated to reproduce the precalculated results of another disorder predictor, Metapredict hybrid, for 363,265 protein sequences from 21 proteomes in order to increase its speed. Performing a similar task, e.g., training a neural network on prepredicted protein sequences from ADOPT, could be an opportunity to increase the speed of our predictor. Both on a single NVIDIA DGX and an Apple M1 chip, I/O time represents ∼ 15% of the inference time and compute time constitutes the remaining ∼ 85%.

We believe that these aspects - accuracy and ease of use - will not just appeal to a wider audience, but also facilitate disorder related research projects that were previously not possible due of scale.

## DISCUSSION

### Remarks on the residue level representations produced by ESM-transformers

There is a set of questions that naturally and frequently arise in the domain of NLP and recently in language models in biological applications: *How much information is encoded in these representations? What exactly do they encode? Are bigger representations always better? In general, how do we make sense of them?*.

To motivate these questions, recall that ODiNPred uses 157 hand-engineered biophysical features as inputs in its prediction model (39). By design, these input features are straightforward to interpret and there is sound rationale as to why they may help predict disorder. In contrast, the input of our ESM-based predictor is much bigger, 1280, which is the size of the representation vectors given by e.g. *ESM-1b*, and its features are, albeit real numbers, abstract, and they are hard to make sense of in physical terms, e.g. residue interaction entropy or net charge.

Answer to the questions above also depend on the downstream task the representations are used for, e.g. predicting fluorescence, contact maps, disorder, sequence/amino acid level classification, etc. This implies a dependency on the prediction model too. In the case of disorder prediction, we use Lasso regression, which is a linear method (56). It is a continuous feature selection technique that means that varying the *shrinkage* parameter *λ* will influence the number of non-zero coefficients. The higher the *λ* is, the greater the *L*_1_ penalty term is and hence the number of non-zero coefficients decreases. In a fitted model with a given *λ*, only those features are used that have non-zero coefficients. In other words, these features are the ones that carry significant information. Therefore, Lasso can help us understand, *which coordinates of the representation vectors are relevant*.

Are all the 1280 coordinates, e.g. in the case of *ESM-1b* needed? We used two different approaches to address this question. The first one was a *naive selection*, whereby only those features are considered, whose coefficients were above a certain threshold in absolute terms. This analysis showed that e.g. with 166 coordinates a correlation of *ρ*_*Spearman*_ = 0.644 could be achieved, as show on Figure 5.

**Figure 5.**
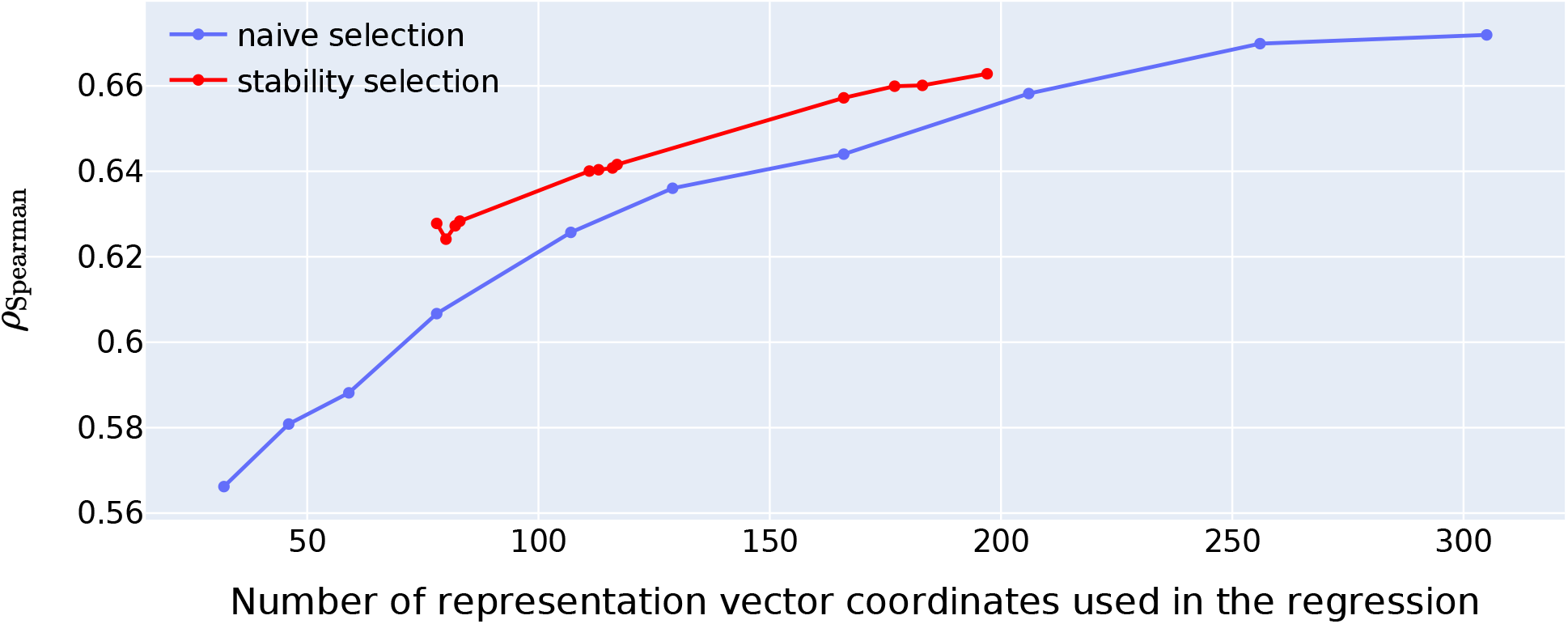
Predictive power of the residue level representation vectors as a function of the number of their coordinates used in the regression. Here, the transformer *ESM-1b* was used, and the predictive power was quantified in terms of *ρ*_Spearman_ between actual and predicted Z-scores on the validation set containing 117 sequences. *Naive selection* (**blue** line) is based on the magnitude of the regression coefficients and *stability selection* (**red** line) is a more robust technique based on resampling; more details are provided in the DISCUSSION section.

**Figure 6.**
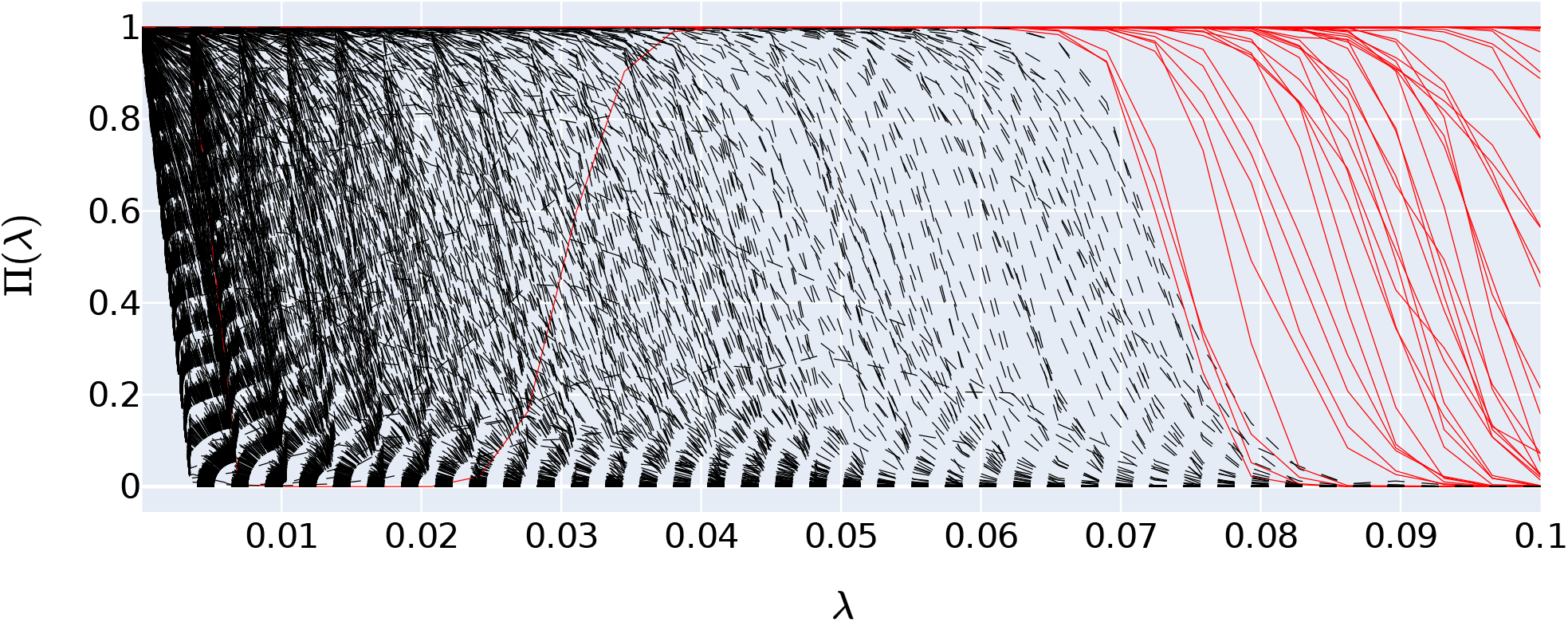
Stability paths examples given by *stability selection*, as described in the MATERIALS AND METHODS section, see Feature Selection subsection. Here, we used the transformer *ESM-1b* and applied the probability and frequency cutoffs *c*_*p*_ = 0.9 and *c*_*f*_ = 20, respectively. A particular line represents one coordinate or feature of the representation vector and shows the selection probability Π_*i*_(*λ*) (on the *y*-axis) as a function of the shrinkage parameter *λ* (on the *x*-axis). The red, solid lines represent the coordinates that made the final subset selection, and the black, dashed lines represent all remaining coordinates. In this particular example, there are 78 red lines. Interestingly, there is one red line (coordinate) buried in the black lines in the region of lower shrinkage values *λ*. This means that even though, more coordinates were selected in Lasso for a lower *λ*, this particular coordinate, which typically was picked for higher *λ*, was not selected.

A better, yet computationally more demanding approach is *stability selection*, which uses resampling, and in this aspect is similar to bootstrapping (58). Using this more robust technique we found that choosing only 83 coordinates of the representation vectors and using those to predict Z-scores already showed an acceptable correlation *ρ*_Spearman_ = 0.6283 on the validation set containing 117 sequences. The predictive power of the representation vectors could be further enhanced by taking additional coordinates, e.g. 117, which yielded a correlation *ρ*_Spearman_ = 0.6416, as depicted by the markers on the red line in Figure 5. Interestingly, this method identified 166 coordinates, which gave a correlation of *ρ*_Spearman_ = 0.6572, which is markedly better than the subset of the same size selected by the naive approach. In comparison to ODiNPred that relies on 157 biophysical features and achieves a correlation of *ρ*_Spearman_ = 0.649 (39), already a subset of 117 coordinates of the *ESM-1b* representations could offer a very similar performance.

This analysis answers, at least partly, the question, whether bigger representations are necessarily better. The *ESM-MSA* based predictor, which uses smaller sized representations, 768, performs relatively poorly with a correlation *ρ*_Spearman_ = 0.6151, see Table 2, compared to all of the alternatives based on the reduced *ESM-1b* representations presented above. This clearly suggests that, at least regarding disorder prediction, the quality of the representations given by *ESM-1b* is genuinely better not purely because of the size of the representation. This is because, we could select a subset of coordinates from the *ESM-1b* representations that was much smaller in size than the size of *ESM-MSA* representations (768), and yet had better prediction power than that of *ESM-MSA*.

### Remarks on the current state of protein disorder predictions

Protein disorder prediction is a challenging task. Even with the methodological advancements of ADOPT, Spearman rank correlation coefficients of *ρ*_Spearman_ up to 0.7 demonstrate that there is still a significant room for further improvements. Beside their different architecture, the most performant protein disorder predictors use deep neural networks to predict the disorder of a residue within a sequence. While these tools identify the features important for disorder prediction, they do not necessarily give a direct indication about the connection of these features to the biophysical origin of protein disorder. Here, we compared two state-of-the-art predictors, ODiNPred (39) and our new ESM-transformer, and found several similarities in their ability to predict certain biophysical features. For example, we could show that both predictors could accurately separate ordered from disordered residues, whose definition is based on the simple observation that experimental Z-scores follow a bimodal distribution. While such a classification performance is promising, applying the predictors to a more diverse dataset, as e.g. used in CAID(34), is more challenging.

We observed that both predictors predict slightly too ordered Z scores. Interestingly, this observation is independent of the actual Z score, e.g., ordered and disordered regions are both predicted, on average, to be too ordered. This indicates that there might be a general feature which contributes to the “orderness” of a residue, which is not completely learned by either of the predictors. Our amino acid analysis reveals that especially the Z-scores of these amino acids, which are known as breakers or stoppers of ordered regions like *α* helices or *β* sheets, e.g. glycine and proline, are less accurately predicted than the other amino acids (Figures 4 and Supplementary Figure S2 and S3 provided in APPENDIX B in the Supplementary Material). This suggests that the prediction of protein disorder might be enhanced if the predictors are trained on well labeled data of these residues as the occurrence of them within a protein sequence plays an important role in the formation of ordered regions.

Furthermore, we observe that it is particularly difficult to predict cysteine residues. Long-range contacts formed from disulfide bridges hold proteins together and are therefore of tremendous importance for the formation of the hydrophobic core of folded proteins. This observation is in line with the analysis of the 157 biophysical features from ODiNPred (39). They observed that the most important features for protein disorder prediction are hydrophobic clusters, the predicted secondary structure, and the evolutionary relationship. While the latter was used to extract the features in this study, it is interesting to see that the other two biophysical features have been learned implicitly by our ESM-transformer, even though it does not use any biophysical features during its training. This shows that the evolutionary information in pre-trained datasets like UniRef(51) already contains these features and transformers are able to unveil this information.

Finally, negligible differences in terms of key performance metrics described in the reliability study under the RESULTS section, clearly illustrate the efficacy of ADOPT in terms on reliability and accuracy.

## CONCLUSIONS

Understanding and accurately predicting protein disorder have been an area of active research in computational biology for more than two decades. Here, we present a sequence based approach to predict disorder using ESM-transformers. These NLP-based methods applied to vast protein databases, e.g. UniProt90, which at the time of writing this paper consisted of 135,301,051 sequences (see https://www.uniprot.org/uniref/. These transformers produce amino acid level representations of protein sequences that we use as inputs in our disorder predictors. We show in various tasks (predicting Z-scores, classifying residues in order/disorder classes) that our predictors offer superior performance compared to the previous state-of-the-art disorder predictor (39) and per-residue scores correlations to protein disorder of models produced by AlphaFold2 (48). We study the quality of these residue level representations by analysing, which coordinates of the residue level representations are relevant in terms of predictive power. We highlight two main advantages of our proposed approach. First, it does not require additional feature engineering or selection of potentially relevant biophysical inputs. Second, its inference capabilities are remarkably fast compared to other publicly available tools. This could aid large scale studies that were previously not possible.

In a broader context, our work complements the most recent developments in NLP based computational biology, see e.g. (68, 69). The common theme of such approaches can be summarized in two steps. First, the aim is to find suitable *embeddings* or *representations* of protein sequences. The second aspect is to use these in relevant *downstream* tasks, e.g. contact map prediction, stability prediction, etc. Downstream use cases are typically less data intense, and this approach is expected to work already reasonably well with data sets of size around 10k or above. Note that in the CheZoD training set alone, which was used in this paper, there are more than 140k observations, given that disorder is a residue level property of interest. In contrast, large transformer models ideally require millions of data points to achieve their full potential. Note that the data points used by transformers are not necessarily experimental observations, but they are purely sequences.

There are many directions for future research related to this particular approach, however, we would highlight three of them specifically. Clearly, great potential lies in the exploration of other relevant downstream tasks. Reiterating the point made in (69), a promising way to improve NLP based techniques, in particular, transformers, in protein science, is to utilise priors, e.g. known protein structures, into these models. This could be a key to enhance the quality of sequence representations. The third area requires novel ideas, but could be invaluable is to connect large transformers to Molecular Dynamics (MD) methods in order to improve simulations and the quality of computed biophysical features. Atomic MD simulations could be used to sample the heterogeneous ensemble of structures in intrinsically disordered regions. The distribution of NMR chemical shifts in these ensembles is directly related to the Z-score and can be used to improve the data used to train transformers while transformers can be used to improve MD force fields.

## Notes

1. https://freesasa.github.io/
2. https://github.com/facebookresearch/esm
3. https://scikit-learn.org/stable/
4. https://github.com/jessevig/bertviz
5. https://github.com/soedinglab/hh-suite
6. https://onnx.ai/
7. We note that these results have been reproduced by Ilzhoefer et al.(70) after the original release of our work in a bioRxiv manuscript from May 26, 2022
8. https://www.uniprot.org/proteomes/UP000005640

## Supplementary Material

## 1 APPENDIX A

### 1.1 Transformer

The Transformer encoder model of ESM is a multi-layer bidirectional Transformer encoder architecture derived from the original implementation[1]. The ESM uses Bidirectional Encoder Representations from BERT-like architecture [2, 3], which alleviates the undirectionality constraint related to a left-to-right architecture where every token can only attend to previous tokens in the self-attention layers of the Transformer. The ESM utilizes a masked language model [4] in which some of the tokens from the input are randomly masked with the objective of predicting the original vocabulary id of the masked residue based only on its context.

#### 1.1.1 Positional encoding

Here **x** is mapped into input the embedding matrix 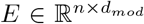, where *E*_*i*,*_ is the embedding vector 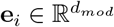 of *x*_*i*_ for *i* = 1, 2, …, *n*.

The vector **e**_*i*_ is defined by the affine transformation, also called linear projection, **e**_*i*_ = **o**_*i*_*W*^*T*^ + **b** ^1^, where *W*^*T*^ indicates the transpose of *W*, **o**_*i*_ ∈ {0, 1} ^*v*^ is the *one-hot encoded* representation of *x*_*i*_ so that *v* = |*V*| is the cardinality of the set *V* while 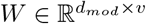 and 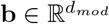 are respectively the weight matrix and the bias computed during the training procedure. The embedding layer is defined as *E* = *embedding*(**x**).

Since the Transformer contains no recurrence and no convolution, in order for the model to make use of the order of the sequence, one must inject some information about the relative or absolute position of the amino acid residues in the protein sequence. To this end, a “positional encodings” **c**_*i*_ is added to the input embedding **e**_*i*_. The ESM makes use of learned positional embeddings [3] i.e an embedding layer fed with the position of each residue in place of the residue itself.

Finally, stacking **d**_*i*_ = **e**_*i*_ + **c**_*i*_ for *i* = 1, 2, …, *n* one gets the matrix 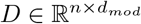.

#### 1.1.2 Attention mechanism

After applying a layer normalization [5], once gets *N*_D_ = *layerNorm*(*D*), where *layerNorm*(*M*) stands for the *layerNorm function* applied on each column of the matrix *M* and 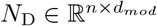.

Three different linear projections are then applied to *N*_D_ getting respectively, except for the bias term, the *query* matrix 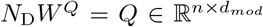, the *key* matrix 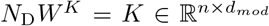 and the *value* matrix 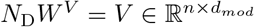 with *W*^*Q*^, *W*^*K*^and 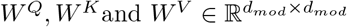 as weight matrices, respectively, computed at training time.

Each of the matrices *Q, K*and *V* is then reshaped into *h* different matrices i.e. 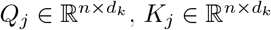 and 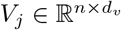 where *j* = 1, 2, …, *h* ∈ ℕ^+^ and 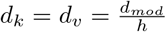.

The *j*-th matrices *Q*_*j*_, *K*_*j*_and *V*_*j*_ are then fed into the *scaled dot-product attention* layer

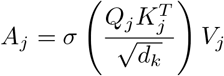

where *σ*(*M*) stands for the *softmax function* [6] applied on each column of the matrix *M*. The matrix *A*_*j*_ is called *attention head j* with 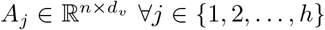 and *h* represents the number of attention heads employed.

The attention heads are then concatenated into *A* = (*A*_1_, *A*_2_, …, *A*_*h*_) with 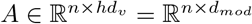 and a linear projection *A*_MH_ = *AW*^*O*^ is applied where, 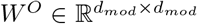 and 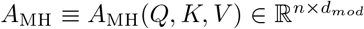. The set of operations which return *A*_MH_ with a matrix *M* as input, is called *multiHeadAttention*(*M*).

A *residual connection* [7] is then employed, getting *R* = *identity*(*N*_D_ + *A*_MH_) where *identity*(*M*) stands for the *identity operator* ^2^ applied on the matrix *M* and 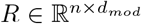. See Supplementary Figure S1 for a multi-head attention output visualisation, related to the CheZod dataset.

**Supplementary Figure S1:**
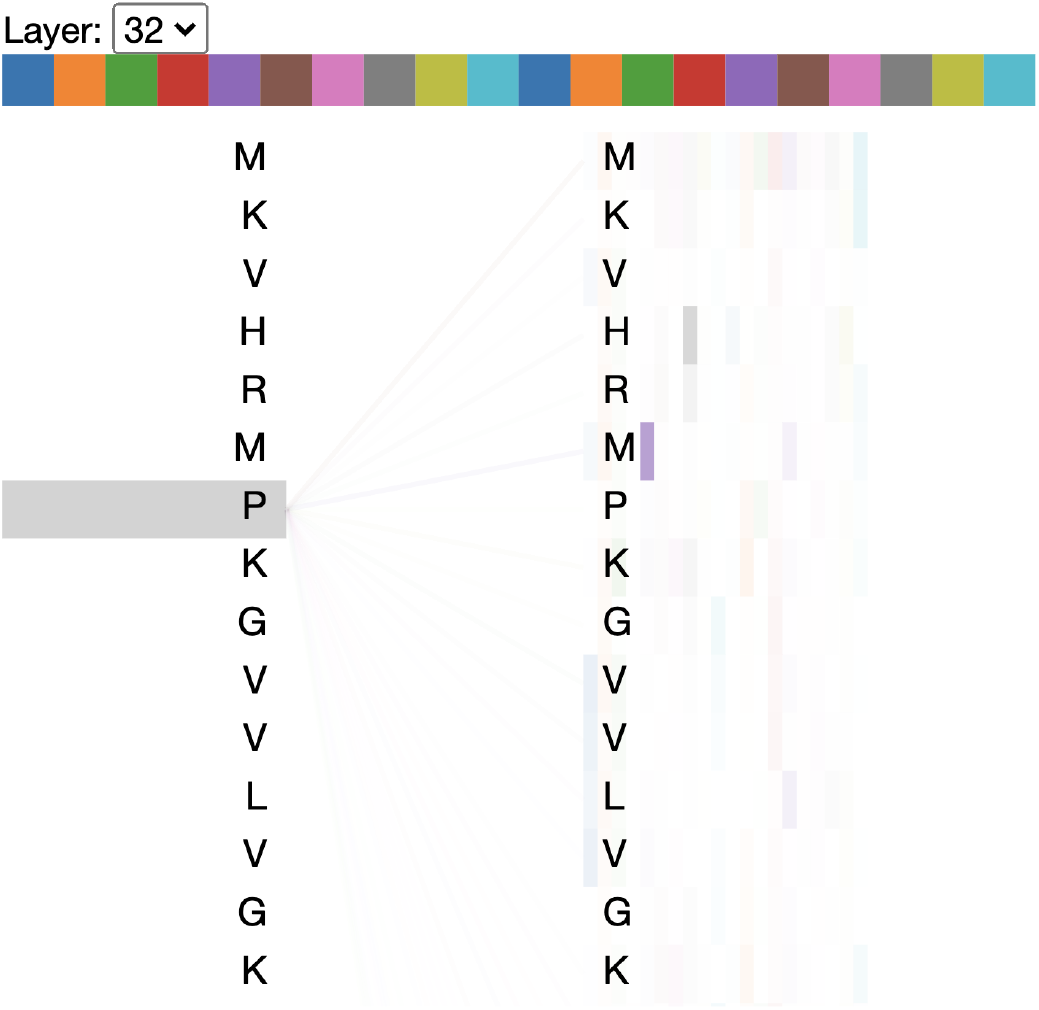
Last multi-attention layer in the **ESM-1b** Transformer model related to a singular protein entry in the CheZod “1325”, identified with the 25096 index. Lines depict the attention from each token (left) to every other token (right). Darker lines indicate higher attention weights, whereas the colours denote different attention heads. The “Layer” drop-down indicates the model layer (zero-indexed). We cut the sequence for visualisation constraints.

#### 1.1.3 Feed-forward network

A *layer normalization* is then applied to the multi-head attention output, so that one gets *N* = *layerNorm*(*R*) with 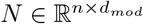.

The matrix *N* is then fed into the *position independent feed-forward network*

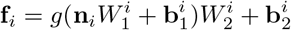

where **n**_*i*_ ≡ *N*_*i*,*_ and *g*(**m**) is the *Gaussian error linear unit* activation function [8] applied to a vector 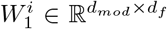 and 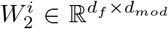 are the weight matrices whereas 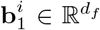 and 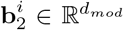 are the bias terms ∀*i* ∈ {1, 2, …, *n*}. Here *d*_*f*_ = 4*d*_*mod*_ for convenience while stacking the position independent feed-forward networks **f**_*i*_ for *i* = 1, 2, …, *n* one gets 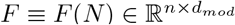. The set of operations which return *F* with a matrix *M* as input is called, *feedForwardNetwork*(*M*).

Finally, *mutatis mutandis*, a *residual connection* is employed, getting *R*_F_ = *identity*(*N* + *F*) with 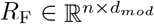. Note that the ESM makes use of *pre-activation blocks*, where the layer normalization is applied prior to the activation and no *dropout* [9] is used.

#### 1.1.4 Encoder block

Once the *positional encoding* layer is applied you get the residue level representation *D* which is then fed into the a *layer normalization* followed by the *multi-head attention* to which a *residual connection* is applied, yielding the residual matrix *R*. The matrix *R* is then fed into another *layer normalization* followed by the *position independent feed-forward network* to which another *residual connection* is applied, yielding *R*_F_.

Therefore the *encoder block* layer can be defined as

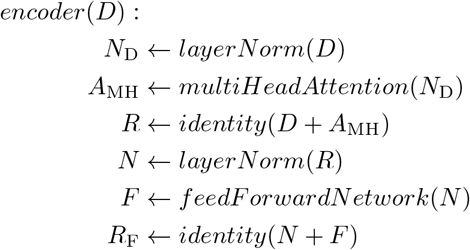

Note that the residue level representation, input matrix *D* of the *encoder block* has the same dimension of the output matrix *R*_F_ of the same block.

#### 1.1.5 Loss

The Transformer has been trained using the *masked language modeling* objective [3] where the input **x** is corrupted by replacing a fraction of the residues with a special mask token “`<mask>`” and the expectation 𝔼 is computed on the set of masked indices at first, and then on the set of protein sequences. The network has been trained to predict the missing tokens from the corrupted sequence, which results in the *minimization* of:

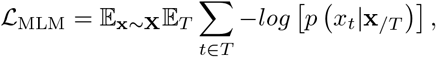

where, for each sequence **x** extracted from the vector of *random variables* **X**, a set of indices *T* is sampled to mask, replacing the true residue *x*_*t*_ with the mask token whereas **x**_*/T*_ represents the masked sequence i.e the context of *x*_*t*_.

The method implemented in [2] has a *token dropout* scheme which replaces the mask token embedding with a fixed vector of zeros so that **e**_*t*_ = **0** ∀*t* ∈ *T* with 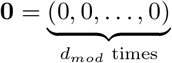.

#### 1.1.3 Masking

The masking strategy in [2] has been adopted from BERT [3], where 15% of the input tokens were selected and predicted through the minimisation of ℒ_MLM_. Of these 80% were replaced with mask token and 10% with a random residue extracted from a uniform distribution; 10% not changed.

#### 1.1.7 Architecture

The Transformer is composed of *l* ∈ ℕ^+^ stacked *encoder blocks*, each fed with the output of the previous one and a final *layer normalization* applied to the output of the last layer.

The output of the last block is denoted as 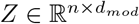 with **z**_*i*_ = *Z*_*i*,*_ and,

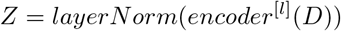

where, 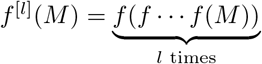.

Please, refer to [2] for additional details.

It is also noteworthy that a projection^3^ back to the size of the vocabulary *v* has been applied to *Z* in order to get the log probabilities one needs to compute ℒ_MLM_.

Finally, the Transformer was trained in batches of *b* ∈ ℕ^+^ proteins, each. Therefore the input was a 2nd rank tensor *X*^*α,μ*^ and each layer was applied on the batch whereas the output was a 3rd rank tensor *Z*^*α,μ,ν*^ where *α* ∈ {1, 2, …, *b*}, *μ* ∈ {1, 2, …, *n*} and *ν* ∈ {1, 2 …, *d*_*mod*_}.

## 2 APPENDIX B

**Supplementary Figure S2:**
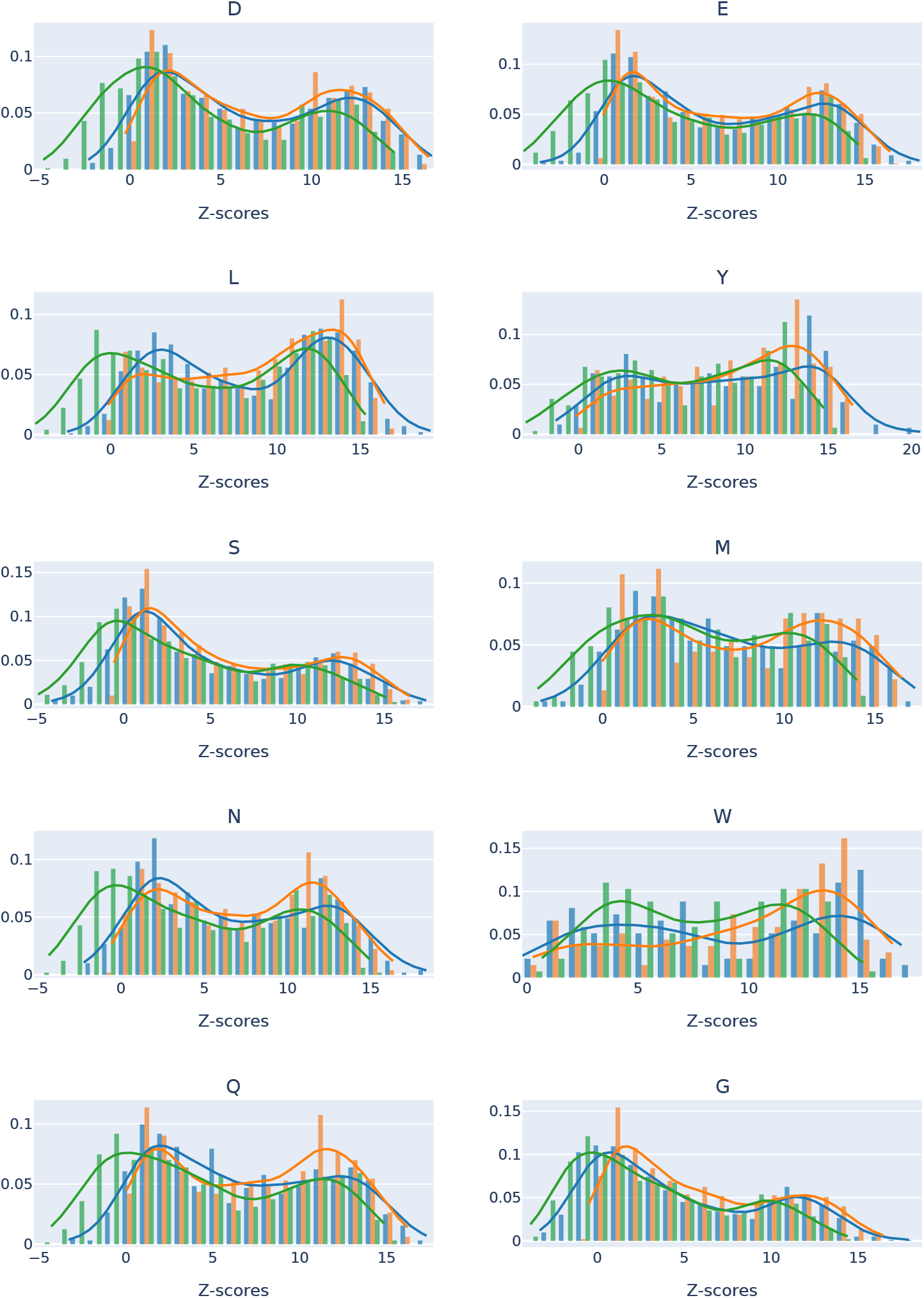
Residue level histograms/density plots of *actual* (**green**), *ESM-1b* (**blue**) and *ODiNPred* (**orange**) predicted Z-scores for residues (from top left to bottom right) [’D’, ’E’, ’L’, ’Y’, ’S’, ’M’, ’N’, ’W’, ’Q’, ’G’]. While in general the ESM transformer based predictor is similar to ODiNPred, in some instances (e.g. residues ’L’ and ’G’) it puts more mass closer to the actual density, than ODiNPred.

**Supplementary Figure S3:**
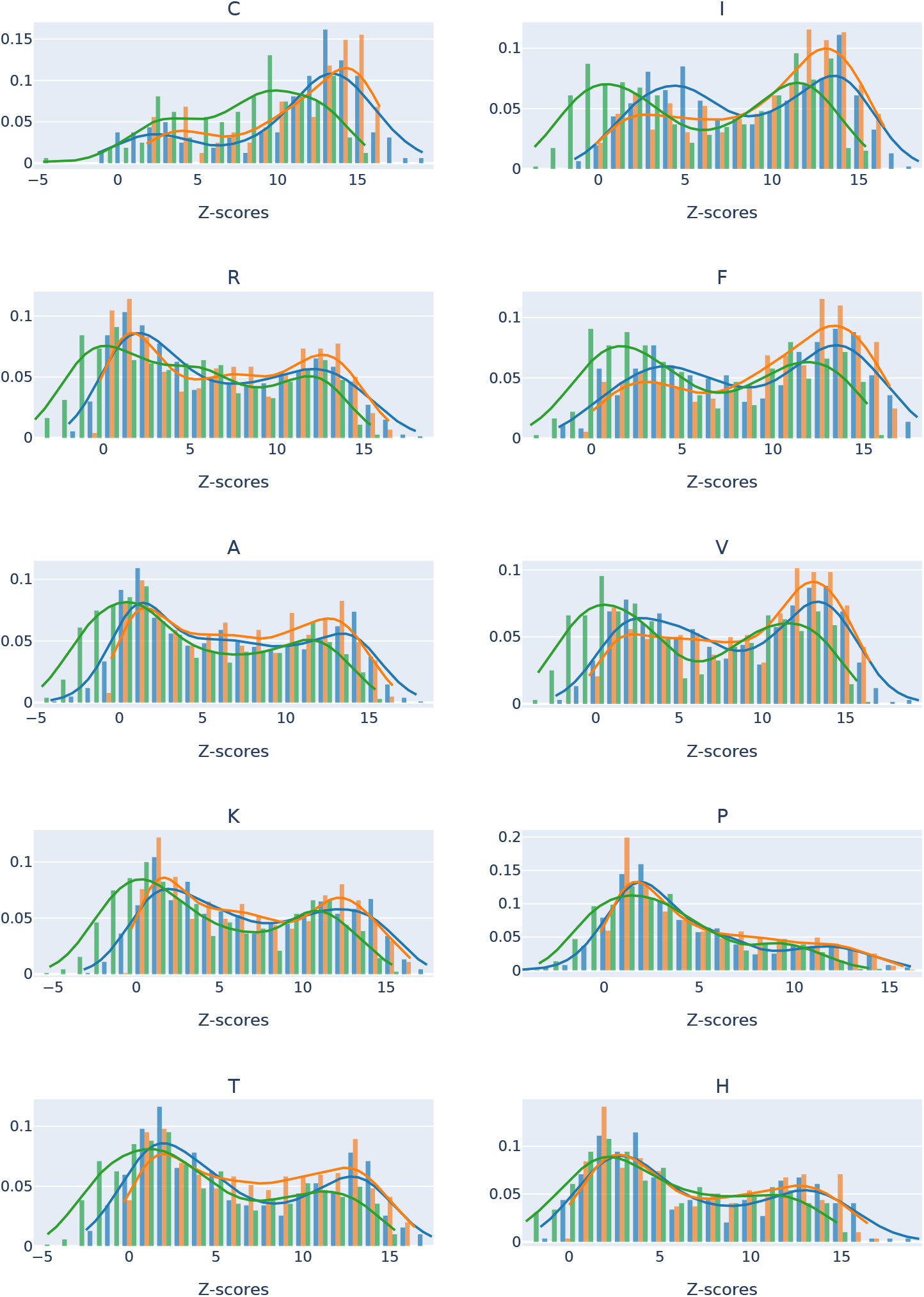
Residue level histograms/density plots of *actual* (**green**), *ESM-1b* (**blue**) and *ODiNPred* (**orange**) predicted Z-scores for residues (from top left to bottom right) [’C’, ’I’, ’R’, ’F’, ’A’, ’V’, ’K’, ’P’, ’T’, ’H’]. While in general the ESM transformer based predictor is similar to ODiNPred, in some instances (e.g. residues ’T’ and ’I’) it puts more mass closer to the actual density, than ODiNPred.

## Notes

1. The linear projection applied of the protein sequence *p* can be represented, except for the bias term, as *D* = *OW*^*T*^ where *O* ∈ ℝ^*n×v*^ is the one hot encoding matrix
2. The identity operator 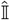 is defined as 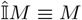
3. A Gaussian error linear unit function and a layer normalisation are applied before the output

## REFERENCES

1. C. B. Anfinsen, “Principles that govern the folding of protein chains,” Science, vol. 181, no. 4096, pp. 223–230, 1973.

2. P. E. Wright and H. Dyson, “Intrinsically unstructured proteins: reassessing the protein structure-function paradigm,” Journal of Molecular Biology, vol. 293, no. 2, pp. 321–331, 1999.

3. P. E. Wright and H. J. Dyson, “Intrinsically disordered proteins in cellular signalling and regulation,” Nature Reviews Molecular Cell Biology, vol. 16, pp. 18–29, 2015.

4. P. Santofimia-Castaño, B. Rizzuti, Y. Xia, O. Abian, L. Peng, Velázquez-Campoy, J. L. Neira, and J. Iovanna, “Targeting intrinsically disordered proteins involved in cancer,” Cellular and Molecular Life Sciences, vol. 77, pp. 1695–1707, 2020.

5. Z. Du and V. N. Uversky, “A comprehensive survey of the roles of highly disordered proteins in type 2 diabetes,” International Journal of Molecular Sciences, vol. 18, no. 10, 2017.

6. Y. Cheng, T. LeGall, C. J. Oldfield, A. K. Dunker, and V. N. Uversky, “Abundance of intrinsic disorder in protein associated with cardiovascular disease,” Biochemistry, vol. 45, pp. 10448–10460, 9 2006. doi: 10.1021/bi060981d.

7. T. P. J. Knowles, M. Vendruscolo, and C. M. Dobson, “The amyloid state and its association with protein misfolding diseases,” Nature Reviews Molecular Cell Biology, vol. 15, pp. 384–396, 2014.

8. G. Fuertes, L. Nevola, and S. Esteban-Martín, “Chapter 9 - perspectives on drug discovery strategies based on idps,” in Intrinsically Disordered Proteins (N. Salvi, ed.), pp. 275–327, Academic Press, 2019.

9. P. Romero, Z. Obradovic, C. Kissinger, J. Villafranca, and A. Dunker, “Identifying disordered regions in proteins from amino acid sequence,” in Proceedings of International Conference on Neural Networks (ICNN’97), vol. 1, pp. 90–95 vol.1, 1997.

10. R. Linding, R. B. Russell, V. Neduva, and T. J. Gibson, “GlobPlot: exploring protein sequences for globularity and disorder,” Nucleic Acids Research, vol. 31, no. 13, pp. 3701–3708, 2003.

11. Z. Dosztányi, V. Csizmok, P. Tompa, and I. Simon, “IUPred: web server for the prediction of intrinsically unstructured regions of proteins based on estimated energy content,” Bioinformatics, vol. 21, no. 16, pp. 3433– 3434, 2005.

12. Z. Dosztányi, V. Csizmók, P. Tompa, and I. Simon, “The Pairwise Energy Content Estimated from Amino Acid Composition Discriminates between Folded and Intrinsically Unstructured Proteins,” Journal of Molecular Biology, vol. 347, no. 4, pp. 827–839, 2005.

13. J. Prilusky, C. E. Felder, T. Zeev-Ben-Mordehai, E. H. Rydberg, O. Man, J. S. Beckmann, I. Silman, and J. L. Sussman, “FoldIndex©: a simple tool to predict whether a given protein sequence is intrinsically unfolded,” Bioinformatics, vol. 21, no. 16, pp. 3435–3438, 2005.

14. O. V. Galzitskaya, S. O. Garbuzynskiy, and M. Y. Lobanov, “FoldUnfold: web server for the prediction of disordered regions in protein chain,” Bioinformatics, vol. 22, no. 23, pp. 2948–2949, 2006.

15. Schlessinger, M. Punta, and B. Rost, “Natively unstructured regions in proteins identified from contact predictions,” Bioinformatics, vol. 23, no. 18, pp. 2376–2384, 2007.

16. J. Cheng, M. J. Sweredoski, and P. Baldi, “Accurate Prediction of Protein Disordered Regions by Mining Protein Structure Data,” Data Mining and Knowledge Discovery, vol. 11, no. 3, pp. 213–222, 2005.

17. K. Peng, P. Radivojac, S. Vucetic, A. K. Dunker, and Z. Obradovic, “Length-dependent prediction of protein intrinsic disorder,” BMC Bioinformatics, vol. 7, no. 1, p. 208, 2006.

18. J. Hecker, J. Y. Yang, and J. Cheng, “Protein disorder prediction at multiple levels of sensitivity and specificity,” BMC Genomics, vol. 9, no. Suppl 1, p. S9, 2008.

19. L. Wang and U. H. Sauer, “OnD-CRF: predicting order and disorder in proteins conditional random fields,” Bioinformatics, vol. 24, no. 11, pp. 1401–1402, 2008.

20. T. Zhang, E. Faraggi, B. Xue, A. K. Dunker, V. N. Uversky, and Y. Zhou, “SPINE-D: Accurate Prediction of Short and Long Disordered Regions by a Single Neural-Network Based Method,” Journal of Biomolecular Structure and Dynamics, vol. 29, no. 4, pp. 799–813, 2012.

21. Walsh, A. J. M. Martin, T. D. Domenico, and S. C. E. Tosatto, “ESpritz: accurate and fast prediction of protein disorder,” Bioinformatics, vol. 28, no. 4, pp. 503–509, 2012.

22. V. Receveur-Bréchot, J. Bourhis, V. N. Uversky, B. Canard, and S. Longhi, “Assessing protein disorder and induced folding,” Proteins: Structure, Function, and Bioinformatics, vol. 62, no. 1, pp. 24–45, 2006.

23. S. Iqbal and M. T. Hoque, “DisPredict: A Predictor of Disordered Protein Using Optimized RBF Kernel,” PLoS ONE, vol. 10, no. 10, p. e0141551, 2015.

24. S. Wang, S. Weng, J. Ma, and Q. Tang, “DeepCNF-D: Predicting Protein Order/Disorder Regions by Weighted Deep Convolutional Neural Fields,” International Journal of Molecular Sciences, vol. 16, no. 8, pp. 17315– 17330, 2015.

25. Hanson, Y. Yang, K. Paliwal, and Y. Zhou, “Improving protein disorder prediction by deep bidirectional long short-term memory recurrent neural networks,” Bioinformatics, vol. 33, no. 5, pp. 685–692, 2016.

26. S. Wang, J. Ma, and J. Xu, “AUCpreD: proteome-level protein disorder prediction by AUC-maximized deep convolutional neural fields,” Bioinformatics, vol. 32, no. 17, pp. i672–i679, 2016.

27. Hanson, K. K. Paliwal, T. Litfin, and Y. Zhou, “SPOT-Disorder2: Improved Protein Intrinsic Disorder Prediction by Ensembled Deep Learning,” Genomics, Proteomics & Bioinformatics, vol. 17, no. 6, pp. 645–656, 2019.

28. C. Mirabello and B. Wallner, “rawMSA: End-to-end Deep Learning using raw Multiple Sequence Alignments,” PLoS ONE, vol. 14, no. 8, p. e0220182, 2019.

29. G. Erdős and Z. Dosztányi, “Analyzing Protein Disorder with IUPred2A,” Current Protocols in Bioinformatics, vol. 70, no. 1, p. e99, 2020.

30. G. Hu, A. Katuwawala, K. Wang, Z. Wu, S. Ghadermarzi, J. Gao, and L. Kurgan, “flDPnn: Accurate intrinsic disorder prediction with putative propensities of disorder functions,” Nature Communications, vol. 12, no. 1, p. 4438, 2021.

31. T. Ishida and K. Kinoshita, “PrDOS: prediction of disordered protein regions from amino acid sequence,” Nucleic Acids Research, vol. 35, no. Web Server issue, pp. W460–W464, 2007.

32. X. Deng, J. Eickholt, and J. Cheng, “PreDisorder: ab initio sequence-based prediction of protein disordered regions,” BMC Bioinformatics, vol. 10, no. 1, pp. 436–436, 2009.

33. P. Kozlowski and J. M. Bujnicki, “MetaDisorder: a meta-server for the prediction of intrinsic disorder in proteins,” BMC Bioinformatics, vol. 13, no. 1, pp. 111–111, 2012.

34. Necci, D. Piovesan, M. T. Hoque, I. Walsh, S. Iqbal, M. Vendruscolo, P. Sormanni, C. Wang, D. Raimondi, R. Sharma, et al., “Critical assessment of protein intrinsic disorder prediction,” Nature Methods, vol. 18, pp. 472–481, 2021.

35. Hatos, B. Hajdu-Soltész, A. M. Monzon, N. Palopoli, L. Á lvarez, B. Aykac-Fas, C. Bassot, G. I. Beńitez, M. Bevilacqua, A. Chasapi, et al., “DisProt: intrinsic protein disorder annotation in 2020,” Nucleic Acids Research, vol. 48, pp. D269–D276, 11 2019.

36. J. T. Nielsen and F. A. A. Mulder, “There is diversity in disorder—”in all chaos there is a cosmos, in all disorder a secret order”,” Frontiers in Molecular Biosciences, vol. 3, p. 4, 2016.

37. E. L. Ulrich, H. Akutsu, J. F. Doreleijers, Y. Harano, Y. E. Ioannidis, J. Lin, M. Livny, S. Mading, D. Maziuk, Z. Miller, et al., “BioMagResBank,” Nucleic Acids Research, vol. 36, pp. D402–D408, 11 2007.

38. K. Tamiola, B. Acar, and F. A. A. Mulder, “Sequence-Specific Random Coil Chemical Shifts of Intrinsically Disordered Proteins,” Journal of the American Chemical Society, vol. 132, no. 51, pp. 18000–18003, 2010.

39. R. Dass, F. A. A. Mulder, and J. T. Nielsen, “ODiNPred: comprehensive prediction of protein order and disorder,” Scientific Reports, vol. 10, p. 14780, Sept. 2020.

40. J. Hanson, Y. Yang, K. Paliwal, and Y. Zhou, “Improving protein disorder prediction by deep bidirectional long short-term memory recurrent neural networks,” Bioinformatics, vol. 33, pp. 685–692, 12 2016.

41. J. Hanson, K. K. Paliwal, T. Litfin, and Y. Zhou, “Spot-disorder2: Improved protein intrinsic disorder prediction by ensembled deep learning,” Genomics, Proteomics & Bioinformatics, vol. 17, no. 6, pp. 645–656, 2019.

42. G. Hu, A. Katuwawala, K. Wang, Z. Wu, S. Ghadermarzi, J. Gao, and L. Kurgan, “fldpnn: Accurate intrinsic disorder prediction with putative propensities of disorder functions,” Nature Communications, vol. 12, p. 4438, 2021.

43. Chatzigeorgiou, V. Constantoudis, F. Diakonos, K. Karamanos, C. Papadimitriou, M. Kalimeri, and H. Papageorgiou, “Multifractal correlations in natural language written texts: Effects of language family and long word statistics,” Physica A: Statistical Mechanics and its Applications, vol. 469, pp. 173–182, 2017.

44. Vaswani, N. Shazeer, N. Parmar, J. Uszkoreit, L. Jones, A. N. Gomez, L. Kaiser, and I. Polosukhin, “Attention is All you Need,” in Advances in Neural Information Processing Systems (I. Guyon, U. V. Luxburg, S. Bengio, H. Wallach, R. Fergus, S. Vishwanathan, and R. Garnett, eds.), vol. 30, Curran Associates, Inc., 2017.

45. J. Devlin, M.-W. Chang, K. Lee, and K. Toutanova, “Bert: Pre-training of deep bidirectional transformers for language understanding,” arXiv preprint arXiv:1810.04805, 2018.

46. Radford, J. Wu, R. Child, D. Luan, D. Amodei, and I. Sutskever, “Language models are unsupervised multitask learners,” OpenAI blog, vol. 1, no. 8, p. 9, 2019.

47. Rives, J. Meier, T. Sercu, S. Goyal, Z. Lin, J. Liu, D. Guo, M. Ott, C. L. Zitnick, J. Ma, et al., “Biological structure and function emerge from scaling unsupervised learning to 250 million protein sequences,” Proceedings of the National Academy of Sciences, vol. 118, no. 15, 2021.

48. J. Jumper, R. Evans, A. Pritzel, T. Green, M. Figurnov, O. Ronneberger, K. Tunyasuvunakool, R. Bates, A. Žídek, A. Potapenko, et al., “Highly accurate protein structure prediction with alphafold,” Nature, vol. 596, pp. 583–589, 2021.

49. J. T. Nielsen and F. A. Mulder, “There is diversity in disorder—”in all chaos there is a cosmos, in all disorder a secret order”,” Frontiers in molecular biosciences, vol. 3, p. 4, 2016.

50. D. S. Wishart, B. D. Sykes, and F. M. Richards, “The chemical shift index: a fast and simple method for the assignment of protein secondary structure through nmr spectroscopy,” Biochemistry, vol. 31, no. 6, pp. 1647–1651, 1992.

51. E. Suzek, Y. Wang, H. Huang, P. B. McGarvey, C. H. Wu, and U. Consortium, “Uniref clusters: a comprehensive and scalable alternative for improving sequence similarity searches,” Bioinformatics, vol. 31, no. 6, pp. 926–932, 2015.

52. Bairoch, R. Apweiler, C. H. Wu, W. C. Barker, B. Boeckmann, S. Ferro, E. Gasteiger, H. Huang, R. Lopez, M. Magrane, et al., “The universal protein resource (uniprot),” Nucleic acids research, vol. 33, no. uppl 1, pp. D154–D159, 2005.

53. M. Mirdita, L. von den Driesch, C. Galiez, M. J. Martin, J. Söding, and M. Steinegger, “Uniclust databases of clustered and deeply annotated protein sequences and alignments,” Nucleic acids research, vol. 45, no. D1, pp. D170–D176, 2017.

54. M. Steinegger, M. Meier, M. Mirdita, H. Vöhringer, S. J. Haunsberger, and J. Söding, “Hh-suite3 for fast remote homology detection and deep protein annotation,” BMC bioinformatics, vol. 20, no. 1, pp. 1–15, 2019.

55. R. Rao, J. Liu, R. Verkuil, J. Meier, J. F. Canny, P. Abbeel, T. Sercu, and Rives, “Msa transformer,” bioRxiv, 2021.

56. T. Hastie, R. Tibshirani, and J. Friedman, The elements of statistical learning: data mining, inference and prediction. Springer, 2 ed., 2009.

57. M. Mirdita, M. Steinegger, F. Breitwieser, J. Söding, and E. Levy Karin, “Fast and sensitive taxonomic assignment to metagenomic contigs,” Bioinformatics, vol. 37, pp. 3029–3031, 03 2021.

58. Meinshausen and P. Bühlmann, “Stability selection,” Journal of the Royal Statistical Society: Series B (Statistical Methodology), vol. 72, no. 4, pp. 417–473, 2010. eprint: https://rss.onlinelibrary.wiley.com/doi/pdf/10.1111/j.1467-9868.2010.00740.x.

59. R. J. Emenecker, D. Griffith, and A. S. Holehouse, “Metapredict v2: An update to metapredict, a fast, accurate, and easy-to-use predictor of consensus disorder and structure,” bioRxiv, 2022.

60. R. J. Emenecker, D. Griffith, and A. S. Holehouse, “Metapredict: a fast, accurate, and easy-to-use predictor of consensus disorder and structure,” Biophysical Journal, vol. 120, pp. 4312–4319, Oct. 2021. Publisher: Elsevier.

61. F.-X. Theillet, L. Kalmar, P. Tompa, K.-H. Han, P. Selenko, A. K. Dunker, G. W. Daughdrill, and V. N. Uversky, “The alphabet of intrinsic disorder,” Intrinsically Disordered Proteins, vol. 1, no. 1, p. e24360, 2013. PMID: 28516008.

62. M. J. Feige, I. Braakman, and L. M. Hendershot, “Chapter 1.1 disulfide bonds in protein folding and stability,” in Oxidative Folding of Proteins: Basic Principles, Cellular Regulation and Engineering, pp. 1–33, The Royal Society of Chemistry, 2018.

63. V. Mariani, M. Biasini, A. Barbato, and T. Schwede, “lDDT: a local superposition-free score for comparing protein structures and models using distance difference tests,” Bioinformatics, vol. 29, pp. 2722–2728, 08 2013.

64. M. Akdel, D. E. V. Pires, E. Porta Pardo, J. Jänes, A. O. Zalevsky, B. Mészáros, P. Bryant, L. L. Good, R. A. Laskowski, G. Pozzati, et al., “A structural biology community assessment of alphafold 2 applications,” bioRxiv, 2021.

65. K. Tunyasuvunakool, J. Adler, Z. Wu, T. Green, M. Zielinski, A. Žídek, Bridgland, A. Cowie, C. Meyer, A. Laydon, et al., “Highly accurate protein structure prediction for the human proteome,” Nature, vol. 596, no. 7873, pp. 590–596, 2021.

66. Piovesan, A. M. Monzon, and S. C. E. Tosatto, “Intrinsic protein disorder and conditional folding in alphafolddb,” Protein Science, vol. 31, no. 11, p. e4466, 2022.

67. J. Wilson, W.-Y. Choy, and M. Karttunen, “Alphafold2: A role for disordered protein/region prediction?,” International Journal of Molecular Sciences, vol. 23, no. 9, 2022.

68. R. Rao, N. Bhattacharya, N. Thomas, Y. Duan, X. Chen, J. Canny, P. Abbeel, and Y. S. Song, “Evaluating Protein Transfer Learning with TAPE,” Advances in neural information processing systems, vol. 32, pp. 9689–9701, Dec. 2019.

69. T. Bepler and B. Berger, “Learning the protein language: Evolution, structure, and function,” Cell Systems, vol. 12, no. 6, pp. 654–669.e3, 2021.

70. Ilzhoefer, M. Heinzinger, and B. Rost, “SETH predicts nuances of residue disorder from protein embeddings,” bioRxiv, p. 2022.06.23.497276, 2022.

## References

[1] A. Vaswani, N. Shazeer, N. Parmar, J. Uszkoreit, L. Jones, A. N. Gomez, L. Kaiser, and I. Polosukhin, “Attention is All you Need,” in Advances in Neural Information Processing Systems (I. Guyon, U. V. Luxburg, S. Bengio, H. Wallach, R. Fergus, S. Vishwanathan, and R. Garnett, eds.), vol. 30, Curran Associates, Inc., 2017.

[2] A. Rives, J. Meier, T. Sercu, S. Goyal, Z. Lin, J. Liu, D. Guo, M. Ott, C. L. Zitnick, J. Ma, et al., “Biological structure and function emerge from scaling unsupervised learning to 250 million protein sequences,” Proceedings of the National Academy of Sciences, vol. 118, no. 15, 2021.

[3] J. Devlin, M.-W. Chang, K. Lee, and K. Toutanova, “Bert: Pre-training of deep bidirectional transformers for language understanding,” arXiv preprint arXiv:1810.04805, 2018.

[4] W. L. Taylor, ““cloze procedure”: A new tool for measuring readability,” Journalism quarterly, vol. 30, no. 4, pp. 415–433, 1953.

[5] J. L. Ba, J. R. Kiros, and G. E. Hinton, “Layer normalization,” arXiv preprint arXiv:1607.06450, 2016.

[6] I. Goodfellow, Y. Bengio, and A. Courville, Deep learning. MIT press, 2016.

[7] K. He, X. Zhang, S. Ren, and J. Sun, “Deep residual learning for image recognition,” in Proceedings of the IEEE conference on computer vision and pattern recognition, pp. 770–778, 2016.

[8] D. Hendrycks and K. Gimpel, “Gaussian error linear units (gelus),” arXiv preprint arXiv:1606.08415, 2016.

[9] N. Srivastava, G. Hinton, A. Krizhevsky, I. Sutskever, and R. Salakhutdinov, “Dropout: a simple way to prevent neural networks from overfitting,” The journal of machine learning research, vol. 15, no. 1, pp. 1929–1958, 2014.

